# Central metabolism as a potential origin of sex differences in morphine analgesia but not in the induction of analgesic tolerance in mice

**DOI:** 10.1101/2020.12.07.414185

**Authors:** Florian Gabel, Volodya Hovhannisyan, Virginie Andry, Yannick Goumon

## Abstract

In rodents, morphine analgesia is influenced by sex. However, conflicting results exist regarding the interaction between sex and morphine analgesic tolerance. Morphine is metabolized in the liver and brain into morphine-3-glucuronide (M3G). Sex differences in morphine metabolism and differential metabolic adaptations during tolerance development might explain the behavioral discrepancies. The present article investigates the differences in peripheral and central morphine metabolism after acute and chronic morphine treatment in male and female mice.

The first experiment aimed to determine whether morphine analgesia and tolerance differ between male and female mice using the tail-immersion test. The second experiment evaluated morphine and M3G metabolic kinetics in the blood using LC-MS/MS. Morphine and M3G were also quantified in several central nervous system (CNS) regions after acute and chronic morphine treatment. Finally, the blood-brain barrier permeability of M3G was assessed in male and female mice.

This study demonstrated that female mice showed weaker morphine analgesia. In addition, tolerance appeared earlier in females but the sex discrepancies observed seemed to be due to the initial differences in morphine analgesia rather than to sex-specific mechanisms involving metabolism. Additionally, compared to male mice, female mice showed higher levels of M3G in the blood and in several CNS regions, whereas lower levels of morphine were observed in these brain regions. These differences are attributable mainly to morphine central metabolism, which differed between males and females in pain-related brain regions, consistent with the weaker analgesic effect in females. However, the role of morphine metabolism in analgesic tolerance seems rather limited.

## INTRODUCTION

Pain management has become one of the most prevalent human health issues with an increasing societal cost. Among painkillers, morphine remains the gold standard to relieve severe pain despite its numerous side effects, including nausea, opioid-induced hyperalgesia (OIH), analgesic tolerance, addiction and ultimately death by respiratory depression (1). Morphine analgesia, as well as the development of its side effects, is influenced by sex in mammals. In rodents, males show more potent analgesia than females with the same dose of morphine (for review, see (2)), whereas human studies have led to more conflicting results (3, 4). Several mechanisms, including hormonal, anatomical, cellular and metabolic disparities, have been proposed to explain these sex differences in animal models (2), although human behavioral discrepancies might also depend on other parameters, such as social context, patient history, strategies used to cope with pain and the presence of comorbidities (5).

Morphine analgesia relies on its binding mainly to μ opioid receptors (MORs) located on neurons of the central nervous system (CNS), especially of brain-regions related to pain such as the lumbar spinal cord (lSC), the periacqueductal gray (PAG) and the amygdala. MORs are also expressed by glial cells, as well as numerous peripheral cells, and mediate various effects, including the modulation of immunity (6). Morphine metabolism involves mainly glucuronidation mediated by the UDP-glucuronosyltransferase (UGT) phase II enzyme family expressed in the liver, intestines, kidneys and, to a significant extent, in some neurons and glial cells (7). In humans, the conjugation of a glucuronide moiety by UGT2B7 on the 3-OH or 6-OH group of morphine produces two predominant metabolites: morphine-3-glucuronide (M3G, 60-70%) and morphine-6-glucuronide (M6G, 10%) (7). In addition, UGT1A1, 1A3, 1A6, 1A8, 1A9, and 1A10 account for minor levels of M3G production (8). However, in mice, UGT2B7 is absent; therefore, no M6G is produced, while most of M3G production is maintained through the action of UGT2B36 (9). M6G has been proposed to be an agonist at MORs, resulting in greater analgesia than morphine (10). In addition, M3G has been described to antagonize morphine effects. Indeed, several studies have reported strong mechanical and thermal hyperalgesia following intraperitoneal, intrathecal, or intracerebroventricular injections of M3G that could block morphine analgesia in rodents (11, 12). Subsequently, many studies have suggested a role of M3G in the development of morphine-induced OIH and analgesic tolerance.

Morphine analgesic tolerance resulting from chronic treatment corresponds to the loss of morphine efficacy and the need for higher doses to achieve sufficient analgesia (13). Although several mechanisms implicating MORs have been previously described to explain this phenomenon (for review, see (14)), neuroinflammatory processes have been proposed to be involved in tolerance mechanisms (15). Interestingly, M3G has been recently shown to elicit pain probably through binding to the Toll-like receptor 4 (TLR4)/myeloid differentiation protein-2 (MD2) complex located on microglial cells and some neurons (11, 16). Consequently, implications of TLR4 activation in analgesic tolerance to morphine and OIH have been described (15, 17). However, conflicting results have argued in opposite directions and correlated OIH and/or tolerance to the MOR rather than to TLR4 (18, 19).

Taken together, numerous pieces of evidence suggest that morphine and M3G have opposing effects. Therefore, the metabolic balance between these two compounds in the periphery and the CNS might govern the analgesic effect of morphine in acute and chronic conditions in males and females. The present article investigates the differences in the metabolic balance in the periphery and the CNS of male and female mice following the acute and chronic, *i.e.*, leading to analgesic tolerance, administration of morphine.

## RESULTS

### Morphine analgesic effect and tolerance in males and females

The tail-immersion test was used to assess sex differences in the analgesic effect of morphine and in the development of morphine analgesic tolerance in the C57BL/6J mouse strain. Daily injections of 10 mg/kg morphine or saline solution were performed for 9 successive days (see protocol **Fig. 1a**). Male and female mice were tested before and 30 min after each injection. Nonlinear regression was applied to the data obtained for each animal. Then, the relevant parameters were compared between the groups. Statistical details are presented in ***SI Appendix***, **Table S1**.

**Fig. 1.**
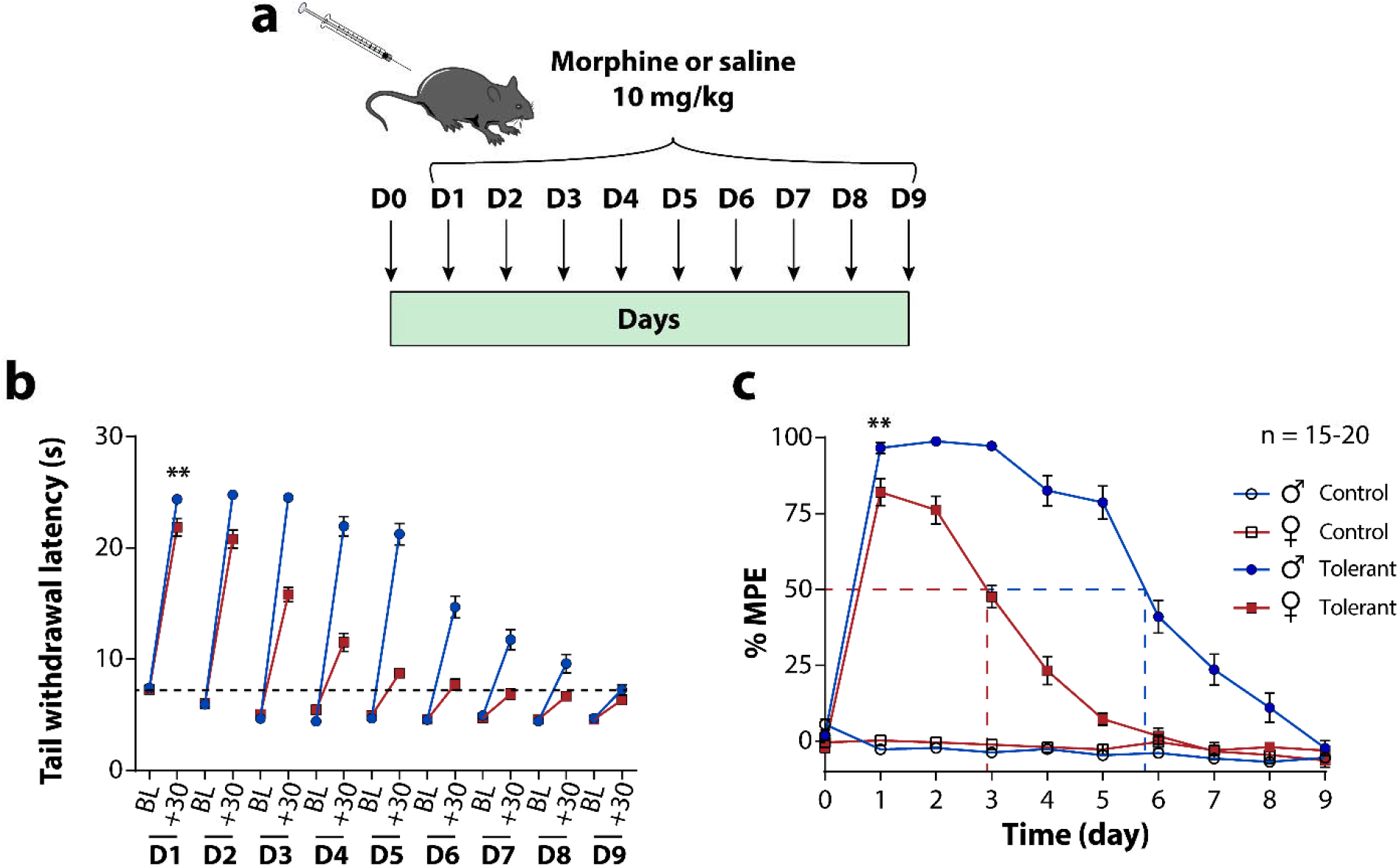
Development of morphine analgesic tolerance in male and female mice. (**a**) Protocol of induction of the analgesic tolerance to morphine. (**b**) Tail withdrawal latencies of male and female mice measured in the tail immersion test before (BL) and 30 min (+30) after morphine injections from day 1 to 9. (**c**) Development of morphine analgesic tolerance throughout the chronic treatment. Anti-nociception is expressed as % of maximum possible effect (% MPE) observed 30 min after morphine or saline injection for 9 successive days. Values are expressed as mean ±SEM; n=15-20 mice per group. Mann-Whitney test was used to compare the analgesic effect of morphine at day 1. **, *P*<0.01. Males are represented as blue circle dots and females as red square dots.

As shown in **Fig. 1b**, the tail-withdrawal latencies measured following the injection of morphine at day 1 were significantly lower in females than in males (Mann-Whitney test, *P*<0.01; **Fig. 1b**). Additionally, this latency decreased over the course of chronic morphine treatment in both males and females but with different kinetics. Moreover, there were no sex differences in the nociceptive threshold of the animals in the basal condition (*i.e.,* before morphine injections). These results showed significant sex differences in the analgesic effect of morphine and in the induction of analgesic tolerance. More precisely, as shown in **Fig. 1c,** the tail-withdrawal latencies were normalized according to the baseline of each animal to visualize the maximal possible effect (MPE) induced by morphine. Female mice showed a morphine MPE of 82.1±4.48% following the first injection of morphine at day 1, which was significantly lower than the MPE of 96.7±1.77% observed in males (Mann-Whitney test, *P*<0.01; **Fig. 1c**). In addition, the morphine MPE decreased with subsequent injections and reached 50% of the MPE on average at day 2.89±0.10 in females and at day 5.88±0.17 in males (unpaired t-test, *P* <0.0001; **Fig. 1c**). Interestingly, no significant difference was observed in the Hill slope coefficient between males (−0.79 ± 0.14) and females (−0.57 ± 0.07), suggesting that the rate of the tolerance development process was identical in males and females.

Moreover, the basal nociceptive threshold (test prior to morphine injections) tended to decrease over the course of the treatment in an identical manner in male and female mice (**Fig. 1b**). This decrease reflected OIH, which seemed to not be influenced by sex.

Together, these results show major sex differences in the analgesic effect of morphine and in the induction of its analgesic tolerance. The differences in tolerance induction appeared to be influenced only by the initial analgesic effects of morphine, which were lower in females than in males.

### Peripheral morphine metabolism

We investigated whether peripheral morphine metabolism differed between males and females following acute and chronic administration of morphine, with the latter leading to analgesic tolerance. At day 10, the concentrations of morphine and M3G in the blood were determined by LC-MS/MS analysis (see protocol **Fig. 2a**). A NCA was separately applied to the data of each animal, and the relevant obtained parameters were compared between the groups with ordinary two-way ANOVA followed by Tukey’s multiple comparisons test. Statistical details are presented in ***SI Appendix,*** **Table S2**.

**Fig. 2.**
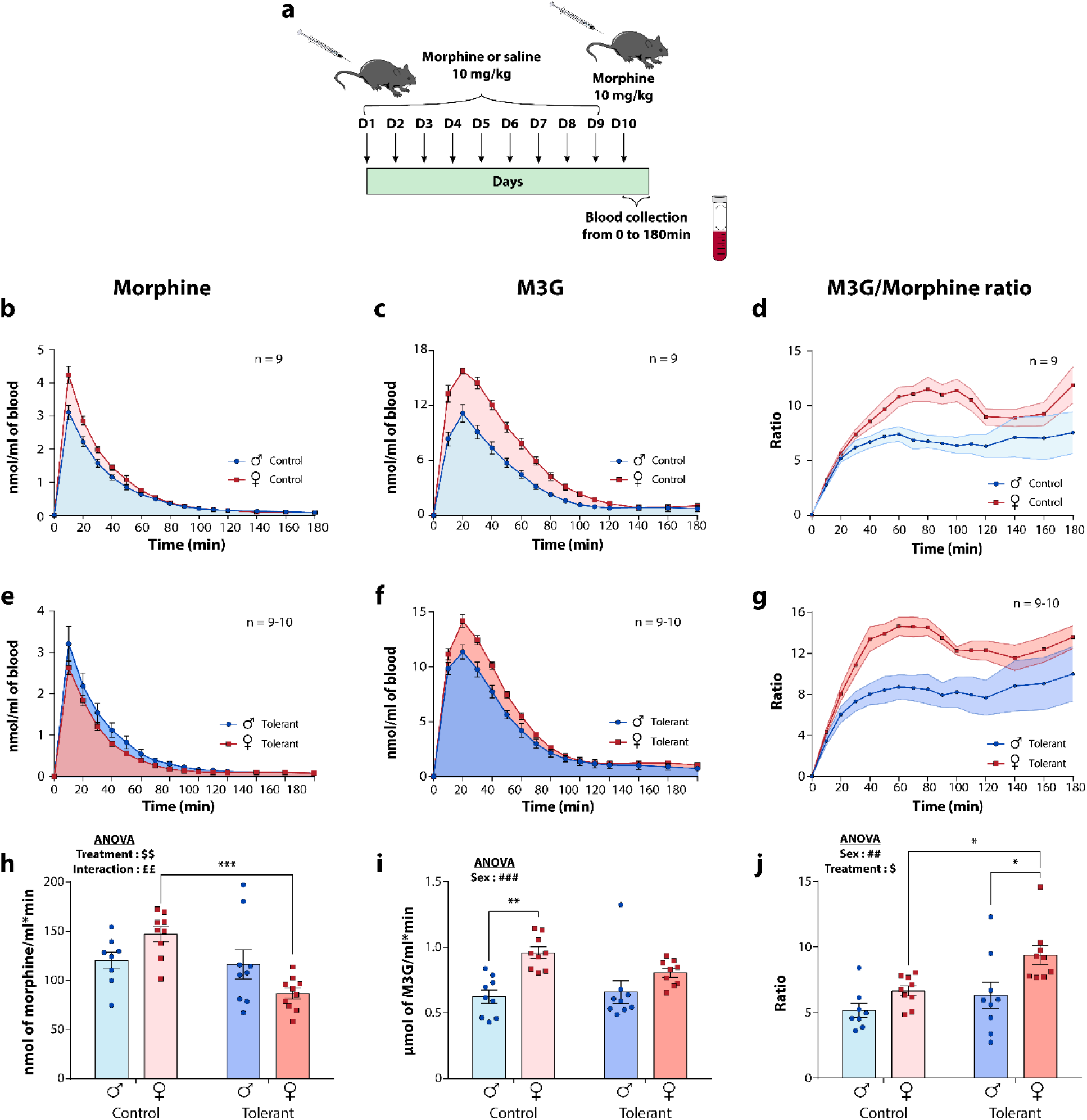
Morphine and M3G kinetics in the blood. (**a**) Protocol of induction of morphine analgesic tolerance across days 1 to 10 (D1-D10, 10 mg/kg morphine or saline i.p.). At day 10, blood was collected at the tail vein at different time points during 180 min. (**b**) Blood levels of morphine in control male and female mice after a single injection of morphine at day 10. (**c**) Blood levels of M3G in control mice. (**d**) M3G/morphine metabolic ratios in the blood of control mice. (**e**) Blood levels of morphine in male and female tolerant mice after an injection of morphine at day 10. (**f**) Blood levels of M3G in tolerant mice. (**g**) M3G/morphine metabolic ratios in the blood of tolerant mice. (**h**) Overall quantities (area under the curve; AUC) of morphine expressed in nmol/ml x min. (**i**) AUC expressed in μmol/ml x min of M3G; (**j**) Ratio M3G/morphine of the corresponding AUC. Values are expressed as means ± SEM, *n =* 9-10. Two-way ANOVA followed by Tukey’s multiple comparisons test was applied. Sex: ##, *P*<0.01; ###, *P*<0.001. Treatment: $, *P*<0.05; $$, *P*<0.01. Interaction: ££, *P*<0.01. *, *P*<0.05; **, *P*<0.01; ***, *P*<0.001. Males are represented as blue circle dots and females as red square dots.

To visualize the sex differences in morphine metabolism, morphine and M3G kinetics and their metabolic ratios over time are depicted in **Fig. 2b, c and d**, respectively, for control mice and in **Fig. 2e, f and g** for tolerant mice. The associated results obtained from the NCA are represented in **Table 1** and as histograms in ***SI Appendix,*** **Fig. S1**.

**Table 1.** Pharmacokinetic parameters obtained from the NCA for morphine and M3G in the blood of male and female control and tolerant mice following an injection of 10 mg/kg of morphine at day 10. Data are expressed as mean ± SEM, *n* = 9-10. Ordinary two-way ANOVA followed by Tukey’s multiple comparisons test was applied. Sex: #, *P*<0.05; ##, *P*<0.01; ###, *P*<0.001; ####, *P*<0.0001. Treatment: $, *P*<0.05; $$, *P*<0.01; $$$, *P*<0.001. Interaction: £, *P*<0.05; ££, *P*<0.01. Control males vs control females: a, *P*<0.05; aa, *P*<0.01; aaaa, *P*<0.0001. Tolerant males vs tolerant females: b, *P*<0.05; bbb, *P*<0.001. Control females vs tolerant females: dd, *P*<0.01, ddd, *P*<0.001; dddd, *P*<0.0001.

|  | **Morphine** |  |  |  |
| --- | --- | --- | --- | --- |
|  | **CT males** | **CT females** | **Tolerant males** | **Tolerant females** |
| **C<sub>max</sub> (nmol/ml)<sup>§, ££</sup>** | 2.93 ± 0.28 | 4.24 ± 0.25<sup>a</sup> | 3.22 ± 0.42 | 2.63 ± 0.16<sup>dd</sup> |
| **AUC<sub>0-inf</sub> (nmol/ml\*min)<sup>\$\$, ££</sup>** | 120.12 ± 8.7 | 146.99 ± 7.7 | 116.41 ± 14.9 | 86.66 ± 52.7<sup>ddd</sup> |
| **AUMC<sub>0-inf</sub> (μmol/ml\*min<sup>2</sup>)<sup>#, \$\$, ££</sup>** | 5.76 ± 0.37 | 6.14 ± 0.36 | 5.80 ± 0.57 | 3.37 ± 0.24<sup>bbb, dddd</sup> |
| **MRT (min)<sup>##</sup>** | 49.10 ± 3.46 | 41.72 ± 1.09 | 52.21 ± 4.63 | 39.06 ± 2.12<sup>b</sup> |
| **T<sub>1/2</sub> (min)<sup>##</sup>** | 34.03 ± 2.40 | 28.91 ± 0.75 | 36.19 ± 3.21 | 27.07 ± 1.47<sup>b</sup> |
| **Cl/F (L/h/kg)<sup>\$\$\$ , £</sup>** | 18.31 ± 1.62 | 14.68 ± 0.91 | 20.25 ± 2.27 | 25.16 ± 1.67<sup>ddd</sup> |
| **Vdss/F (L/kg)<sup>§</sup>** | 15.51 ± 2.61 | 10.18 ± 0.61 | 18.16 ± 2.74 | 16.44 ± 1.46 |
|  | **M3G** |  |  |  |
|  | **CT males** | **CT females** | **Tolerant males** | **Tolerant females** |
| **C<sub>max</sub> (nmol/ml)<sup>####</sup>** | 11.38 ± 0.85 | 16.35 ± 0.39<sup>aaaa</sup> | 11.52 ± 0.72 | 14.21 ± 0.57<sup>b</sup> |
| **AUC<sub>0-inf</sub> (nmol/ml\*min)<sup>###</sup>** | 624.4 ± 50.3 | 959.7 ± 42.9<sup>aa</sup> | 659.1 ± 85.8 | 805.8 ± 32.2 |
| **AUMC<sub>0-inf</sub> (μmol/ml\*min<sup>2</sup>)** | 33.62 ± 4.1 | 47.22 ± 3.2 | 39.14 ± 12.0 | 44.90 ± 5.4 |
| **MRT (min)** | 53.12 ± 3.6 | 48.85 ± 1.4 | 53.22 ± 6.3 | 54.88 ± 4.8 |

#### Effect of sex

The statistical analysis revealed a significant effect of sex on the area under the first moment curve (AUMC; two-way ANOVA, sex: *P*<0.05; **Table 1**), mean residence time (MRT; two-way ANOVA, sex: *P*<0.01; **Table 1**) and half-life (two-way ANOVA, sex: *P*<0.01; **Table 1**) of morphine. Moreover, a strong sex effect was observed in the maximal concentration of M3G that was reached over the time course (two-way ANOVA, sex: *P*<0.0001; **Table 1**), as well as on the M3G area under the curve (AUC; two-way ANOVA, sex: *P*<0.001; **Fig. 2i**). Importantly, a significant effect of sex was thus observed on the metabolic M3G/morphine AUC ratio (two-way ANOVA, sex: *P*<0.01; **Fig. 2j**). There was no impact of sex with any other reported parameters even though a trend was observed in the volume of distribution of morphine at steady state (two-way ANOVA, sex: *P=*0.09; **Table 1**).

#### Effect of treatment

The analysis revealed an effect of the treatment on the maximal concentration of morphine reached over the time course (two-way ANOVA, treatment: *P*<0.05; **Table 1**), morphine AUC (two-way ANOVA, treatment: *P*<0.01; **Fig. 2h**), morphine AUMC (two-way ANOVA, treatment: *P*<0.01; **Table 1**), morphine clearance (two-way ANOVA, treatment: *P*<0.001; **Table 1**) and volume of distribution of morphine at steady state (two-way ANOVA, treatment: *P*<0.01; **Table 1**). In addition, there was no effect of treatment on the reported M3G parameters (**Table 1**). Consequently, a significant effect of the treatment was noted on the metabolic M3G/morphine AUC ratio (two-way ANOVA, treatment: *P*<0.05; **Fig. 2j**).

#### Interaction

Several interactions were observed between the reported parameters. More precisely, the maximal concentration reached over the time course (two-way ANOVA, interaction: *P*<0.01; **Table 1**), AUC (two-way ANOVA, interaction: *P*<0.01; **Fig. 2h**), AUMC (two-way ANOVA, interaction: *P*<0.01; **Table 1**) and clearance (two-way ANOVA, interaction: *P*<0.05; **Table 1**) of morphine were influenced by both variables. In addition, a trend was observed in the M3G maximal concentration reached (two-way ANOVA, interaction: *P=*0.09; **Table 1**). These interactions were mainly driven by the differences between control and tolerant female mice, which were not observed in male mice (see **Table 1**, post hoc analysis). It is thus impossible to make conclusions regarding the main effects with these parameters. Nevertheless, no interaction was seen with the M3G parameters (**Table 1**) or in the metabolic ratios (**Fig. 2j**), suggesting that peripheral morphine metabolism into M3G did not seem to be differentially involved during the development of analgesic tolerance to morphine in male and female mice. These interactions were more likely related to changes in morphine absorption and/or clearance.

Taken together, our results indicated that (***i***) female mice displayed much higher peripheral morphine metabolism and had significantly higher levels of M3G than males, and (***ii***) the peripheral metabolism of morphine was exacerbated during the development of analgesic tolerance to morphine in mice.

### Quantification of morphine and M3G in brain regions and the lumbar spinal cord

On day 10, morphine and M3G levels were quantified by LC-MS/MS in the amygdala, PAG, lSC and OB 30 min after the injection of morphine in control and tolerant male and female mice (see protocol **Fig. 3a)**. The values obtained with morphine and M3G are reported in the table insert in **Fig. 3b** and illustrated as histograms in ***SI Appendix***, **Fig. S2**. Statistical details are presented in ***SI Appendix***, **Table S3**. Ordinary two-way ANOVA was first used to assess the global effect of sex, treatment, and their potential interaction, whereas specific differences between the groups were evaluated with Tukey’s multiple comparisons test. In addition, to account for the sex disparities observed in morphine and M3G levels, the concentrations found in various brain regions, the lSC and the blood of each mouse were normalized (according to those found in males) and are represented in **Fig. 3c and 3d**. In both control (**Fig. 3c**) and tolerant mice (**Fig. 3d**), it clearly appeared that females show overall lower levels of morphine and higher levels of M3G than males.

**Fig. 3.**
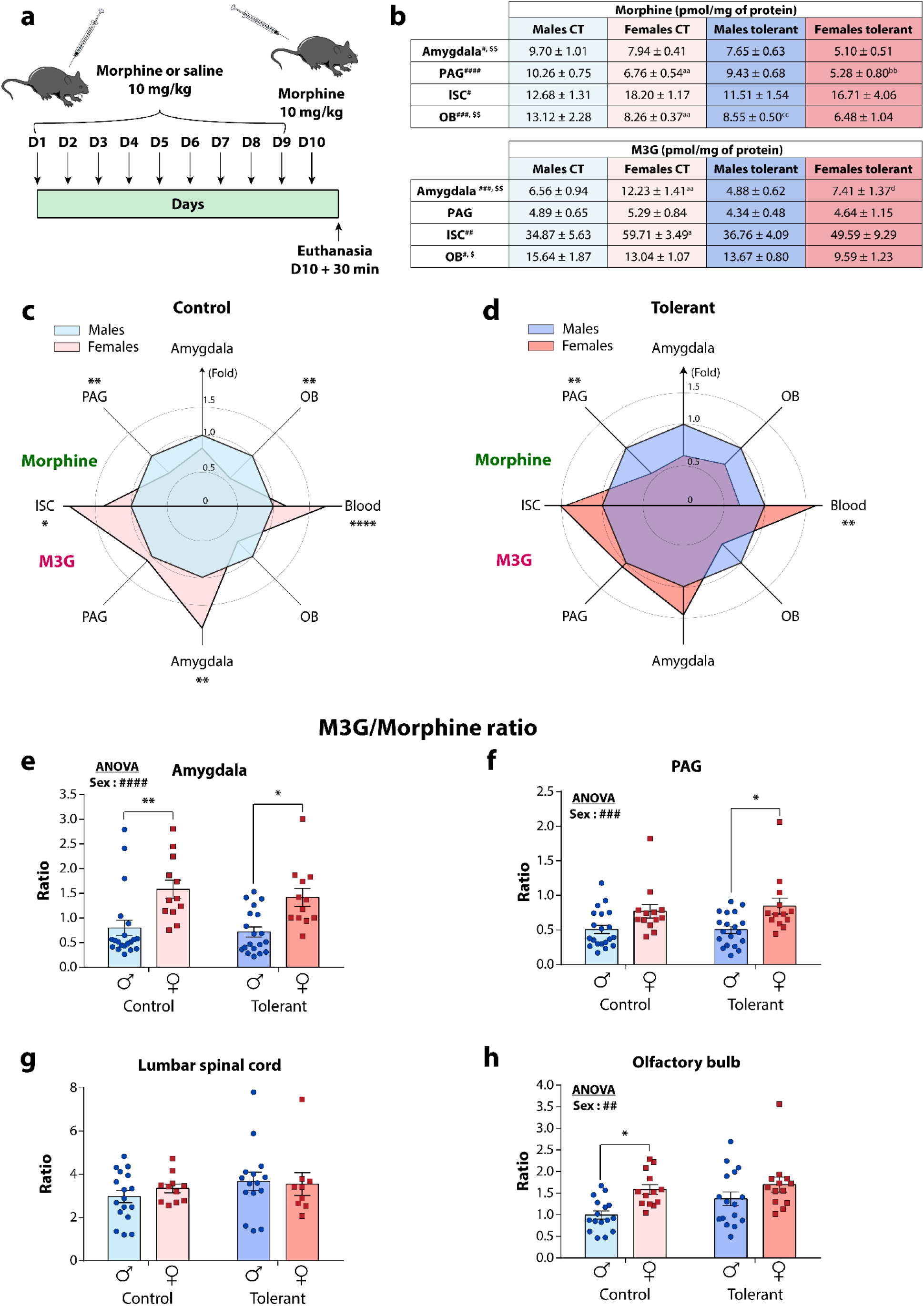
Levels of morphine and M3G in the different brain areas and lumbar spinal cord of male and female control and tolerant mice. (**a**) Protocol of induction of morphine analgesic tolerance across days 1 to 10 (D1-D10, 10 mg/kg of morphine or saline i.p.). At day 10, brain areas and lumbar spinal cord were collected 30 min after the injection of morphine and, morphine and M3G were quantified by LC-MS/MS. (**b**) Levels of morphine and M3G found in the amygdala, the PAG, the ISC and the OB. Morphine and M3G levels in the brain regions, the lSC and the blood of each mouse were normalized according to those found in males in (**c**) control and (**d**) tolerant mice. M3G/morphine ratios found in (**e**) the amygdala, (**f**) PAG, (**g**) lSC and (**h**) OB. Values are expressed as means ± SEM, *n =* 9-20. Ordinary two-way ANOVA followed by Tukey’s multiple comparisons test was applied. Sex: #, *P*<0.05; ##, *P*<0.01; ###, *P*<0.001; ####, *P*<0.0001. Treatment: $, *P*<0.05; $$, *P*<0.01. Control males vs control females: a, *P*<0.05; aa, *P*<0.01. Tolerant males vs tolerant females: bb, *P*<0.01. Control males vs tolerant males: cc, *P*<0.01. Control females vs tolerant females: d, *P*<0.05. Tukey’s multiple comparisons results are reported as *, *P*<0.05; **, *P*<0.01; ****, *P*<0.0001 in (**c**) and (**d**). Males are represented as blue circle dots and females as red square dots.

#### Effect of sex

The analysis revealed that significantly lower levels of morphine were present in the amygdala (two-way ANOVA, sex: *P*<0.01; **Fig. 3b**), PAG (two-way ANOVA, sex: *P*<0.0001; **Fig. 3b**) and OB (two-way ANOVA, sex: *P*<0.001; **Fig. 3b**) of the female mice compared to the male mice. Surprisingly, morphine concentrations were higher in female lSC than in male lSC (two-way ANOVA, sex: *P*<0.05; **Fig. 3b**). Furthermore, a much higher level of M3G was found in the amygdala (two-way ANOVA, sex: *P*<0.001; **Fig. 3b**), lSC (two-way ANOVA, sex: *P*<0.001; **Fig. 3b**) and OB (two-way ANOVA, sex: *P*<0.05; **Fig. 3b**) of female mice. No sex differences in M3G levels were observed in the PAG (**Fig. 3b**). Interestingly, the metabolic ratio between M3G and morphine was notably lower in male mice than in female mice in the amygdala (two-way ANOVA, sex: *P*<0.0001; **Fig. 3e**), PAG (two-way ANOVA, sex: *P*<0.001; **Fig. 3f**)) and OB (two-way ANOVA, sex: *P*<0.01; **Fig. 3h**), whereas sex did not influence the M3G/morphine ratio in the lSC (**Fig. 3g**).

#### Effect of treatment

Two-way ANOVA indicated a significant effect of treatment on morphine levels in the amygdala (two-way ANOVA, treatment: *P*<0.01; **Fig. 3b**) and in the OB (two-way ANOVA, treatment: *P*<0.05; **Fig. 3b**). In addition, an effect of the treatment was also observed on M3G concentrations in the amygdala (two-way ANOVA, treatment: *P*<0.01; **Fig. 3b**) and the OB (two-way ANOVA, treatment: *P*<0.05; **Fig. 3b**). However, there was no effect of the treatment on morphine or M3G levels in the PAG and the lSC. Interestingly, there was no effect of treatment on the M3G/morphine metabolic ratio, although a trend was noticed in the OB (two-way ANOVA, treatment: *P=*0.081; **Fig. 3h**).

Together, these results suggested major discrepancies in morphine and M3G levels, as well as in their metabolic ratio, between males and females. These differences, at least in the amygdala and the PAG, are correlated with the behavioral sexual dimorphism observed in the analgesic effect of morphine. However, the induction of morphine tolerance did not modify the metabolic ratio even though there were differences in morphine and M3G levels in tolerant mice compared to control mice. This suggested a rather limited effect of tolerance on the balance between morphine and M3G in the CNS regions that were analyzed. Furthermore, sex was not implicated in the differences between control and tolerant mice, as witnessed by the absence of any interactions between the two factors.

### Morphine and M3G brain/blood ratios

To investigate the origin of the differences in morphine and M3G levels and metabolic ratios in the different groups of animals, we determined whether these differences (***i***) are the consequence of the differences observed in peripheral metabolism, (***ii***) are due to differences in M3G BBB permeability, and/or (***iii***) are dependent on the central metabolism of morphine.

First, we established brain/blood ratios to normalize the concentrations of morphine or M3G found in the CNS regions based on those found in the blood of each animal. Ordinary two-way ANOVAs followed by Tukey’s multiple comparisons tests were performed to identify the potential differences between groups. Statistical details are presented in ***SI Appendix,*** **Table S4**.

#### Effect of sex

Two-way ANOVA indicated a significant sex effect on morphine brain/blood ratios in the PAG (two-way ANOVA, sex: *P*<0.0001; **Fig. 4d**) and lSC (two-way ANOVA, sex: *P*<0.01; **Fig. 4g**), although in opposite directions. Furthermore, an effect of sex was observed in M3G brain/blood ratios in the OB (two-way ANOVA, sex: *P*<0.0001; **Fig. 4k**) and PAG (two-way ANOVA, sex: *P*<0.05; **Fig.4e**).

**Fig. 4.**
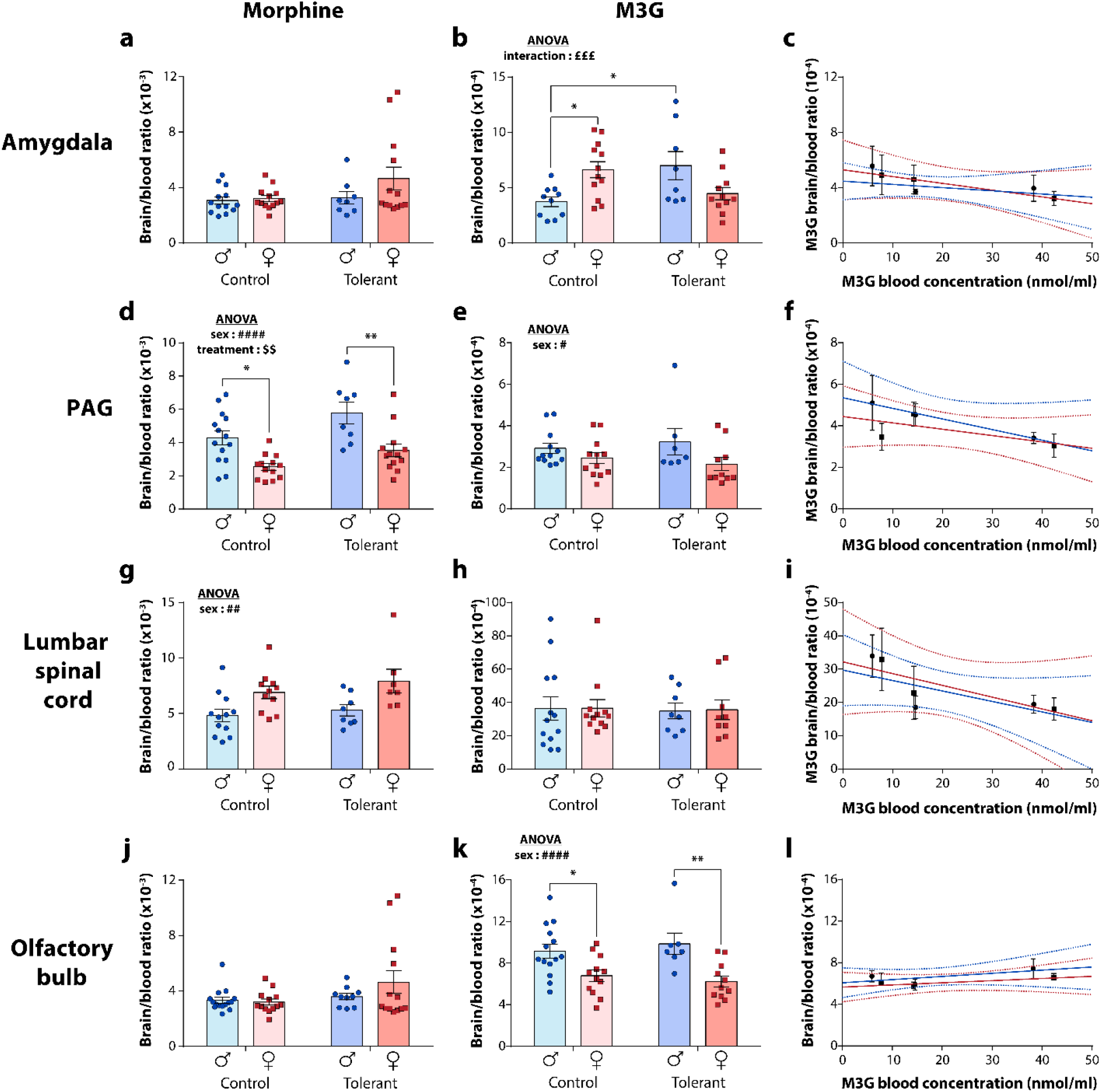
Brain/blood ratio of morphine and M3G in the different brain areas and lumbar spinal cord of male and female control and tolerant mice. Brain/blood ratio of (**a**) morphine and (**b**) M3G in the amygdala. (**c**) M3G brain/blood ratio obtained in the amygdala as a function of M3G concentration found in the blood after i.p. injections of increasing concentrations of M3G. Brain/blood ratio of (**d**) morphine and (**e**) M3G in the PAG. (**f**) M3G brain/blood ratio obtained in the PAG as a function of M3G concentration found in the blood after i.p. injections of increasing concentrations of M3G. Brain/blood ratio of (**g**) morphine and (**h**) M3G in the lSC. (**i**) M3G brain/blood ratio obtained in the lSC as a function of M3G concentration found in the blood after i.p. injections of increasing concentrations of M3G. Brain/blood ratio of (**j**) morphine and (**k**) M3G in the OB. (**l**) M3G brain/blood ratio obtained in the OB as a function of M3G concentration found in the blood after i.p. injections of increasing concentrations of M3G. The blue and red lines represent linear modelling of the BBB permeability for M3G in males and females, respectively. 95% Confidence intervals are represented as dotted-line with the appropriate color. Values are expressed as means ± SEM, *n =* 7-15. Ordinary two-way ANOVA followed by Tukey’s multiple comparisons test was applied. Sex: #, *P*<0.05; ##, *P*<0.01; ####, *P*<0.0001. Treatment: $$, *P*<0.01. Interaction: £££, *P*<0.001. *, *P*<0.05; **, *P*<0.01. Males are represented as circle dots and females as square dots.

#### Effect of treatment

ANOVA showed an unexpected significant impact of the treatment on morphine brain/blood ratios in the PAG (two-way ANOVA, treatment: *P*<0.01; **Fig. 4d)**. Additionally, a trend was observed in the OB (two-way ANOVA, treatment: *P=*0.076; **Fig. 4j**). However, there was no effect of the treatment on M3G brain/blood ratios.

#### Interaction

An interaction between the effects was observed for the M3G brain/blood ratios only in the amygdala (two-way ANOVA, interaction: *P*<0.001; **Fig. 4b**). Interestingly, the post hoc analysis revealed a significant effect between control male and female mice in the amygdala (Tukey’s multiple comparisons test, *P*<0.05; **Fig. 4b**). In addition, a significantly higher M3G brain/blood ratio was found between control and tolerant males in the same structure (Tukey’s multiple comparisons test, *P*<0.05; **Fig. 4b**). This last difference was observed only in males, resulting in the interaction. Further investigations are required to understand its origin.

Taken together, these results suggested that the sex differences in morphine and M3G levels observed in the different CNS regions between males and females do not necessarily reflect the differences found in the blood. Morphine and/or M3G BBB permeability or central metabolism of morphine could be partially responsible for such differences. However, it appeared that the repeated morphine injection protocol had a rather limited influence on the brain/blood ratios, suggesting that the differences in morphine and M3G levels observed in the CNS regions, with the exception of the PAG, might reflect those observed in the blood.

### M3G blood-brain barrier permeability in males and females

As the main differences observed in the M3G brain/blood ratios differed by sex, we evaluated to what extent the BBB permeability for M3G differed between males and females. Different doses of M3G (10, 20 and 40 mg/kg) were injected into naïve male and female mice. After 30 min, the levels of M3G were quantified in the blood, the brain regions of interest and the lSC. Linear regression was used, and the models obtained in the males and females were compared with the extra sum-of-squares F-test. Statistical details are presented in ***SI Appendix,*** **Table S5**.

As shown in **Fig. 4**, there were no differences in M3G BBB permeability in any analyzed structure when the M3G brain/blood ratios were plotted as a function of the M3G blood concentration in female and male mice (red and blue lines, respectively; **Fig. 4c, 4f, 4i, 4l**). In addition, the BBB permeability for M3G seemed to be relatively linear with increasing doses of M3G, suggesting a passive diffusion mechanism (***SI Appendix,*** **Fig. S3**).

### Central metabolism of morphine

Then, we hypothesized that the differences observed in M3G brain/blood ratios in the amygdala, PAG and OB relied on the central metabolism of morphine that differed between male and female animals. Hence, we evaluated whether the M3G levels found in the brain regions of interest after an injection of morphine were significantly different from the M3G levels obtained in the same structure after an injection of M3G based on the M3G concentration found in the blood. Linear regression was applied, and the extra sum-of-squares F-test was used to compare models. Statistical details are presented in ***SI Appendix,*** **Table S5**.

As shown in **Fig. 4**, based on the M3G concentrations found in the blood of each mouse, the M3G brain/blood ratios obtained in the amygdala after the injection of morphine were significantly higher than those obtained after an injection of M3G in female mice (extra sum-of-squares F-test, *P*<0.05; **Fig. 4**) but not in males. In addition, it appeared that female mice show higher M3G brain/blood ratios than male animals (extra sum-of-squares F-test, *P*<0.01; **Fig. 4**). These results indicated that morphine was metabolized into M3G directly in the CNS and that this metabolism differed between male and female mice. In contrast, male mice showed more robust central morphine metabolism in the OB than females (extra sum-of-squares F-test, *P*< 0.01; **Fig. 4**). However, the M3G brain/blood ratios reported in the PAG were unexpectedly low, and there were no differences in the lSC.

Taken together, these results suggested that morphine is metabolized in the CNS *in vivo* in important areas relevant to pain. In addition, important sex differences were observed in the central metabolism of morphine that could explain the behavioral differences observed in the analgesic effects of morphine between male and female animals.

## DISCUSSION

### Sex differences in morphine analgesic effect and metabolism

Our behavioral experiments showed that in female C57Bl/6J mice, 30 min after the injection, morphine displayed a 15% lower analgesic effect compared to that in male mice. This difference in effectiveness was relatively weak compared to the literature, although it is explained by the cutoff of 25 s used in the tail-immersion test (20, 21). Indeed, sex differences in morphine analgesia have been described in both humans and rodents. However, the results from human studies have often been contradictory (2, 22, 23). Alternatively, a vast majority of rodent studies have shown that morphine elicited weaker analgesia in females than in males (2). Nevertheless, there are also discrepancies across rodent studies based on species, genotypes and paradigms used to assess the analgesic effects of morphine (24, 25).

The origin of the disparity in the analgesic effect of morphine between males and females remains controversial. Many mechanisms have been proposed, including organizational and activational differences (26), differential expression of the MOR (27), functional differences in the recruited pain circuit (28), dimorphism in glial cell activation (29), and a potential role for drug metabolism (30). For instance, morphine analgesia has been shown to vary according to the estrous cycle in females (27). However, in our experiment, female mice were not synchronized during the tolerance setting, and our behavioral results show only small variations in the response to morphine in females. Therefore, it seems unlikely that, in our paradigm, the estrous cycle played a major role in the sexual dimorphism observed with morphine analgesia and tolerance.

Interestingly, the morphine and M3G concentrations found in the blood and in the CNS of control mice were consistent with the higher potency of morphine observed in male C57BL/6J mice (20). Female mice showed higher morphine metabolism after a single i.p. injection of morphine, and these results were consistent with the differences observed by South *et al.* in 2009 after intravenous (i.v.) injection (31). Even though a surprising difference was observed for morphine levels in the blood, the dramatically higher M3G concentrations found in the female blood is consistent with the 2-fold higher M3G/morphine ratio observed in females throughout the time course of the monitoring. These results suggest a dramatic imbalance in the analgesic *vs* pronociceptive effects of morphine and M3G, respectively.

Several hypotheses may explain such differences. One hypothesis is that in female mice, morphine is metabolized into M3G at a higher rate than in males. Indeed, sex differences in UGT expression have previously been reported in the literature (32). In addition, we determined a significantly higher morphine half-life (±15 and ±25% in control and tolerant mice, respectively) and MRT (±15 and ±25% in control and tolerant mice, respectively) in male mice than in female mice. However, such a difference alone cannot explain the higher metabolic ratio observed in the blood of female mice.

A second hypothesis is that more of the morphine is converted into M3G than the other morphine metabolites (which usually account for up to 30%) in females (7). However, this hypothesis is unlikely, as the major proportion of morphine is metabolized into M3G in male mice, and a small increase in this proportion in females cannot explain the strong differences observed in the metabolic ratio.

Finally, it is also possible that the differences observed in the metabolic ratio rely on the morphine and M3G distribution in the body and/or its renal excretion. Indeed, Rush *et al.* showed no difference in morphine glucuronidation by hepatic microsomes in male and female rats (33). In addition, sex differences have previously been shown in the distribution of glucuronide metabolites. For instance, Bond *et al,* in 1981, showed that DNBalcG, one of the dinitrotoluene glucuronidated metabolites, is found at higher levels in the bile of male compared to female rats (34). Together, it is possible that the distribution in the body and excretion of morphine and M3G differ between males and females, leading to higher concentrations of M3G in the blood of female mice. Nevertheless, this hypothesis requires further investigation to be validated.

To summarize, exacerbated peripheral morphine metabolism in females, as well as sex differences in the distribution and/or excretion of morphine and M3G, might be responsible for an imbalance between morphine-mediated analgesia and M3G-induced hyperalgesia in females. Moreover, experiments performed mainly on rats have shown that differences in the M3G/morphine plasma ratio might play a role in male-female differences observed in morphine antinociception (24, 35). However, even though sex differences in mouse hepatic metabolism of morphine were observed in our experiment, Sarton *et al.* in 2000 did not observe any sex differences in morphine, M3G and M6G levels in the plasma of healthy volunteers (23). In any case, the contradictory results between humans and mice could be explained by species differences in liver UGT expression (36). In addition, it is improbable that sex-related differences in morphine BBB permeability or hepatic metabolism might fully explain the differences observed in morphine analgesia. In agreement with this statement, Kest *et al.* in 1999 observed sex differences in response to morphine in the tail-flick test following direct i.c.v. injections (20), suggesting that the BBB might not be implicated or only be implicated to a limited extent in sex-related differences in analgesia.

Therefore, our main hypothesis is that the behavioral differences in the analgesic effect of morphine between male and female mice rely on the central metabolism of morphine. Indeed, several *in vitro* studies have shown the capability of brain homogenates and glial cells to metabolize morphine into M3G in both mice and humans (13, 37). In addition, even though M3G displays low BBB permeability (38), we showed here that higher levels of M3G were present in different brain regions following an i.p. injection of morphine compared with after an i.p. injection of M3G, consistent with the data reported for the whole brain of guinea pigs (39).

Taken together, our results suggest that morphine metabolism takes place in some areas of the brain *in vivo*. Interestingly, the M3G brain/blood ratios were higher in females than in male mice, at least in the amygdala. Surprisingly, the opposite result was found in the OB. In addition, we did not observe any differences in BBB permeability for M3G between males and females in any brain regions tested. These results indicate important sex-dependent differences in the central metabolism of morphine *in vivo*. Furthermore, even though the M3G brain/blood ratios observed in the PAG after an injection of morphine were unexpectedly lower than those reported after the injection of M3G, morphine and M3G brain/blood ratios reported in the PAG in male mice were higher than those detected in female mice. These results suggest that (***i***) the central metabolism of morphine takes place in the PAG but does not seem to be influenced by sex and (***ii***) the BBB permeability for morphine is different between males and females.

One should also note that the M3G half-life reported by Handal *et al.* in 2002 after an injection of M3G is approximately 30 min, whereas we reported an MRT for M3G between 45 min and 55 min following an injection of morphine (40). Therefore, the total amount of M3G present in the blood of the animal before quantification in the brain 30 min after injection is likely much higher after administration of M3G than after administration of morphine. Hence, as we show that the BBB permeability of M3G increased proportionally with its blood concentration, the central metabolism of morphine is probably much higher and underestimated in our experiment.

In agreement with these statements, morphine levels found in the PAG and amygdala of male animals were significantly higher than those found in female animals. Morphine has been described as producing potent analgesic effects through its action mainly in the CNS. Furthermore, M3G levels found in the amygdala of female animals were greater than those in male animals, even though the opposite results were observed with morphine. This result is consistent with the lower analgesic effect of morphine observed in females since several studies described a neuroexcitatory and pronociceptive effect of M3G following intrathecal and intracerebroventricular injections (11, 41). Alternatively, Peckmann *et al.*, in 2005 reported higher ED50 in female rats than in males for several opiates that produces 3-glucuronides metabolites (42). Hence, M3G and other 3-glucuronide metabolites might act as excitatory signals, and M3G levels found in the brain might modulate morphine analgesia in mice. However, conflicting results have been reported including studies showing no pronociceptive effects of M3G (43, 44).

Consequently, in our study, the M3G/morphine ratios were strongly increased in the PAG and in the amygdala of female compared to male mice. Importantly, Barjavel *et al,* in 1995, correlated the analgesic effect of morphine following s.c. injection in male rats with the M3G/morphine ratio found in the cortical extracellular fluid in a microdialysis study (45). It is worth noting that we have surprisingly observed contrasting results in the lSC, even though there were no differences in the M3G/morphine ratios.

Taken together, our results indicate that the metabolism of morphine occurs in the brain *in vivo* and is differentially influenced by sex in C57BL/6J mice. This results in a modulation of morphine and M3G levels in some pain-related CNS regions. Thus, sex differences in the central metabolism of morphine, as well as sex differences in the distribution and/or excretion of M3G, might be responsible for a shift in the balance between morphine analgesia and M3G hyperalgesia in females. In addition, the roles for BBB permeability and sex hormones in these sex differences are unlikely. Future studies will investigate to what extent these differences in metabolic balance and distribution contribute to the behavioral contrast observed in the analgesic effect of morphine.

### Metabolism involvement in analgesic tolerance to morphine

We observed strong sex differences in the development of analgesic tolerance to morphine in C57BL/6J mice. This tolerance appeared 3 days earlier in females than in males during the protocol. However, the rate at which this tolerance developed remained the same between males and females, as witnessed by the absence of differences in the Hill slope coefficients. Furthermore, the analgesic effect of morphine at day 1 was significantly lower in females than in males, although the MPE of morphine was reached in males during the first 3 days of the protocol due to the 25-s cutoff set for the tail-immersion test. These results suggest that, in our paradigm, the disparities observed in the development of morphine tolerance are due to differences between males and females in morphine analgesia at day 1 that are likely underestimated rather than to a sex-specific mechanism involved in the development of morphine tolerance. In addition, the development of OIH started immediately on day 2 and was identical in males and females. However, the paradigm used to assess morphine analgesia and tolerance was not optimal to evaluate morphine-related OIH; thus, clear sex differences in OIH might have been difficult to measure.

Additionally, regarding morphine peripheral metabolism, we did observe significant interactions at day 10 between sex and chronic treatment in the maximal concentration, AUC, AUMC and clearance of morphine. All these interactions were based on the lower AUC in tolerant female mice compared to their respective controls, while this effect was not observed in males. However, there was no interaction in the reported parameters for M3G or on the metabolic ratios obtained in the blood. Moreover, there was no interaction reported in any condition tested in the brain, with the exception of the M3G brain/blood ratios in the amygdala. These results suggest that the rapid induction of analgesic tolerance to morphine in females might not be related to sex-specific mechanisms involving morphine metabolism. However, it should be noted that sexual dimorphism in analgesic tolerance has been previously documented, although strong discrepancies were noticed regarding the species and paradigms used to assess morphine tolerance (25, 46).

We observed that tolerant mice had lower levels of morphine in the blood than control animals. In addition, the metabolic ratios between M3G and morphine were increased in the blood of tolerant mice, suggesting that chronic morphine injections exacerbated the metabolism of morphine. Moreover, we observed an increase in morphine volume of distribution at steady state in tolerant mice, suggesting an increase in morphine distribution. Interestingly, the mRNA of UGTs implicated in testosterone metabolism has been shown to be upregulated in the liver following a single morphine administration (47). Therefore, it is possible that morphine can directly or indirectly regulate UGT and transporter expression, hence modulating its own metabolism, distribution and/or excretion. Consequently, lower levels of morphine found in the blood and increased metabolic M3G/morphine ratios might partially be responsible for the decrease in the analgesic effect of morphine. It is, however, highly unlikely that these differences play a major role in morphine tolerance, as tolerant male animals show the same metabolic ratio as control females. This suggests that the alteration in morphine metabolism might be responsible for at most 15 to 30% of the loss of morphine analgesic effects.

Regardless, the quantification in the blood was consistent with the morphine quantification in the amygdala and OB. Decreased levels of morphine were observed in the amygdala and OB in tolerant mice 30 min after injection. However, morphine was still significantly present and should have continued to produce an analgesic effect. Furthermore, M3G levels remained unchanged in every region tested, with the exception of the amygdala and OB where the levels were decreased, eliminating a potential increase in its pronociceptive effect due to higher concentrations in the brain. In addition, we did not observe any changes in the PAG and lSC, as expected in light of their major role in morphine analgesia. Finally, the metabolic ratio between M3G and morphine in the analyzed CNS regions did not differ between control and tolerant mice, excluding a potential role for central metabolism in the development of analgesic tolerance to morphine.

## CONCLUSION

In conclusion, our results showed sex differences in morphine analgesia and metabolism following a single administration in mice. Females displayed lower analgesia following a single administration of morphine consistent with (***i***) greater levels of M3G found in the blood, (***ii***) lower levels of morphine and greater levels of M3G found in some pain-related brain regions. In addition, the differences observed in these brain regions were related to the central metabolism of morphine that occurred to a greater extent in some pain-related brain regions in female mice. Hence, this could be responsible for the sex differences observed in morphine analgesia.

In addition, morphine tolerance appeared earlier during the protocol in females than in males, although the rate of its development seemed to not be influenced by sex. The strong disparities observed in the induction of tolerance were thus due to the existing sex differences in morphine analgesia. In addition, tolerant mice showed lower concentrations of morphine in the blood, as well as higher M3G/morphine metabolic ratios. However, globally, no changes were observed in the brain regions of tolerant mice even though lower levels of morphine were observed in the amygdala and OB. Together, hepatic morphine metabolism was exacerbated by chronic morphine treatment; however, central metabolism did not appear to be involved in morphine analgesic tolerance.

All these data support morphine hepatic and central metabolism as related to sex differences observed in morphine analgesia in C57BL/6J mice. In addition, the role of these factors in analgesic tolerance to morphine seems to be relatively limited.

## MATERIALS AND METHODS

### Animals

Experiments were performed with 10 weeks-old male and female C57BL/6J mice (26±4 g and 20±4 g, respectively; Charles River, L’Arbresle, France). Animals were housed according to a 12 h light-dark cycle, at a temperature of 23°C±2°C and provided with food and water *ad libitum*. All procedures were performed in accordance with European directives (2010/63/EU) and were approved by the regional ethics committee and the French Ministry of Agriculture (license No. APAFIS# 23671-2020010713353847 v5 and APAFIS#16719-2018091211572566 v8 to Y.G.).

### Induction of morphine analgesic tolerance

To evoke morphine analgesic tolerance, mice were weighed and injected intraperitoneally (i.p.) with either 10 mg/kg of morphine (w/v, Francopia, Paris, France) dissolved in NaCl 0.9% or with an equal volume of saline solution every morning (light phase at 10 AM) for 9 consecutive days. On day 10, all mice received an injection of 10 mg/kg of morphine with a calibrated Hamilton syringe before the final procedure.

### Behavioural assessment of morphine analgesic effect

The analgesic effect of morphine was measured with the tail immersion test. Mice were first habituated to their environmental conditions for a week without any experimental procedures. Then, they were gently handled and habituated to be restrained in a grid pocket for two days. Mice were tested every day by measuring the latency of the tail withdrawal when 2/3 of the tail was immersed in a constant-temperature water bath heated at 47°C. In the absence of response, the cut-off was set at 25 s to avoid tissue damage. The basal thermal nociceptive threshold was determined during two weeks of baseline and considered as steady following three consecutive days of stable measurement prior to the testing phase. Mice were tested before and 30 min after the injection of morphine or saline solution for 9 successive days. Results are expressed as % maximal possible effect according to the following formula:

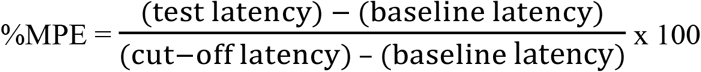

### Blood collection

On day 10, tails of the mice were anaesthetized locally with a topic application of lidocaine/prilocaine 5% (Zentiva, Paris, France). After 5 min, a small incision was performed at the end of the tail and 5 μl of blood were collected using a heparinized calibrated capillary (Minicaps End-to-End 5 μl; Hischmann, Eberstadt Germany). Then, all mice were injected with morphine, and 5 μL of blood were collected every 10 min for 2 hours and every 20 min for the last hour.

### Brain regions and lumbar spinal cord sampling

On day 10, mice were euthanized 30 min following the injection of morphine, and brains were removed and placed on an ice-cold mouse brain matrix. Razor blades were used to cut the brain into 1mm thick slices. Punchers of 1 mm and 0.5 mm diameters were used to sample the periaqueductal gray (PAG) and amygdala, respectively. Olfactory bulbs (OB) were extracted using forceps. For the lumbar spinal cord (lSC), hydraulic extrusion was performed as described before (48). Structures were directly transferred in micro-tubes and stored at −80°C.

### M3G blood-brain barrier permeability

To investigate whether differences in M3G blood-brain barrier (BBB) permeability exist between males and females, 15 male and 15 female mice were weighed, divided into 3 groups and injected i.p. with either 10 mg/kg, 20 mg/kg or 40 mg/kg of M3G (w/v, Sigma Aldrich, St. Quentin Fallavier). Mice were euthanized 30 min following the injection of M3G and the blood, the brain regions of interest and the lSC were collected according to the protocol described above.

### Sample preparation

#### Blood

The blood was transferred from the capillary into a micro-tube containing 4 μl of heparin and frozen at −20°C for later analysis. On the day of the analysis, blood was thawed, and 10 μl of internal standard (IS; containing 12 pmol of D3-morphine and 10.5 pmol of D3-M3G; Sigma Aldrich) and 100 μl of ice-cold acetonitrile (ACN; Thermo Scientific, San Jose, USA) were added. The samples were vortexed and centrifuged at 20,000g during 15 min at 4°C. The supernatants were collected, dried under vacuum and suspended in 800 μl of H_2_O/0.1% formic acid (v/v; Sigma Aldrich) prior to solid-phase extraction (SPE). HyperSep PGC SPE-cartridges (1cc, 25 mg, Thermo Electron, Villebon Sur Yvette, France) were used with a positive pressure manifold (Thermo Electron). Briefly, cartridges were activated with 1 ml of ACN followed by a two-step wash with 2 ml of H_2_O/0.1% formic acid (v/v). Then, samples were loaded onto the cartridges and dried for a minute under high vacuum. The cartridges were subsequently washed with 1 ml of H_2_O/0.1% formic acid (v/v) followed by 1 ml of 97.9% H_2_O/2% ACN/0.1% formic acid (v/v). Elution was performed with 800 μl of 79.9% H_2_O/20% ACN/0.1% formic acid (v/v), and eluates were centrifuged at 20,000g, 4°C for 5 min. Supernatants were dried under vacuum and resuspended in 50 μl of H_2_O/0.1% formic acid (v/v) prior to LC-MS/MS analysis.

#### Brain regions and lumbar spinal cord

Samples were sonicated (2×5 s, 100W) in 200 μl of H_2_O containing 10 μl of IS (containing 40 pmol of D3-morphine and 60 pmol of D3-M3G). After centrifugation for 15 min at 20,000g and 4°C, 10 μl of the supernatants were precipitated with 100 μl of ice-cold ACN for 30 min. Supernatants were dried under vacuum after another centrifugation for 15 min at 20,000g and 4°C and resuspended in 20 μl of H_2_O/0.1% formic acid (v/v) prior to LC-MS/MS analysis.

### LC-MS/MS instrumentation and analytical conditions

Analyses were performed with a Dionex Ultimate 3000 HPLC system (Thermo Electron) coupled with a triple quadrupole Endura mass spectrometer (Thermo Electron). Xcalibur v4.0 software was used to control the system (Thermo Electron). Samples were loaded onto a ZORBAX SB-C18 column (150 x 1 mm, 3.5 μm, flow of 90 μl/min; Agilent, Les Ulis, France) heated at 40°C. LC and MS conditions used are detailed in ***SI Appendix,*** **Table S6**.

Identification of the compounds was based on precursor ions, selective fragment ions and retention times obtained for the heavy counterpart present in the IS. Selection of the monitored transitions and optimization of collision energy and RF Lens parameters were determined manually (for details, see ***SI Appendix,*** **Table S6**). Qualification and quantification were performed using the multiple reaction monitoring mode (MRM) according to the isotopic dilution method (49).

### Non-compartmental analysis

Pharmacokinetic parameters for morphine and M3G were determined through a non-compartmental analysis (NCA) performed with PKsolver described by Zhang *et al.* in 2010 (50). The λ_z_ acceptance criteria were set as followed: R adjusted > 0.80, includes ≥ 3 time points, AUC_tlast-inf_ ≤ 20% AUC_0-inf_. The linear up log down trapezoidal rule was used to determine the AUC of morphine and M3G after extrapolation to infinity.

### Statistics

Statistical analysis was performed using GraphPad Prism 6 Software. All experiments were conducted according to a 2×2 factorial design, and groups were compared using ordinary two-way ANOVA followed by post-hoc Tukey’s multiple comparisons test.

For the behavioral experiments, non-linear regression with a 4-parameters logistic equation was applied to the data to extract the following parameters of each animal: MPE% at day 1, time at which half of the MPE is reached and the Hill slope coefficient. Then, the mean of each parameters was compared using either an unpaired t-test or Mann-Whitney test after a normality check with the D'Agostino & Pearson omnibus normality test.

For the comparison between the M3G brain/blood ratio obtained following an injection of morphine and M3G, linear regressions were applied and analyzed through a nested-model comparison with the extra sum-of-squares F-test.

Results are presented as mean values ± standard error of the mean (SEM). A p-value <0.05 was considered statistically significant.

## ABBREVIATIONS

ACN: acetonitrile
AUC: area under the curve
AUMC: area under the first moment curve
BBB: blood-brain barrier
CID: collision gas
Cl/F: clearance over bioavailability
Cmax: maximal concentration reached over the time course
CNS: central nervous system
d3-morphine: morphine bearing three ^2^H
LC-MS/MS: liquid chromatography coupled to tandem mass spectrometry
lSC: lumbar spinal cord
M3G: morphine-3-glucuronide
M6G: morphine-6-glucuronide
MOR: mu opioid receptor
MPE: maximal possible effect
MRM: multiple reaction monitoring mode
MRT: mean residence time
NCA: non-compartmental analysis
OB: olfactory bulb
OIH: opioid induced hyperalgesia
PAG: periaqueductal gray
T_1/2_: half-life
UGT: UDP-glucuronosyl-transferase
V_dss_/F: volume of distribution at steady-state over bioavailability

## ACKNOWLEDGMENTS AND GRANT SPONSOR

This work was funded by CNRS, University of Strasbourg, French Ministère Délégué à la Recherche et à l'Enseignement Supérieur (PhD fellowship to F.G., and V.O.). We thank the following research programs of excellence for their support: FHU Neurogenycs, French National Research Agency (ANR) through the Programme d'Investissement d'Avenir (contract ANR-17-EURE-0022, EURIDOL graduate school of pain).

**SI Appendix, Table S1.** Statistical details for morphine analgesic effect and induction of tolerance (Figure 1). Non-linear regression with a 4-parameters logistic equation was applied to the data of each animal. Then, the obtained parameters were averaged and compared with an unpaired t-test with Welch’s correction or a Mann-Whitney test according to the results of the D'Agostino & Pearson omnibus normality test. MPE, maximal possible effect.

|  | Time at which half %MPE |  | Hill coefficient |  | %MPE at day 1 |  |
| --- | --- | --- | --- | --- | --- | --- |
|  | D'Agostino & Pearson omnibus normality test |  |  |  |  |  |
|  | Males | Females | Males | Females | Males | Females |
| K2 | 0.94 | 1.04 | 3.89 | 5.90 | 23.9 | 8.03 |
| p-value | 0.63 | 0.59 | 0.14 | 0.052 | <0.0001* | 0.018* |
|  | Unpaired t-test with Welch's correction |  | Unpaired t-test with Welch's correction |  | Mann-Whitney test |  |
| t, df | t=15.40 df=27.33 |  | t=1.419 df=25.31 |  | / |  |
| U | / |  | / |  | 78.0 |  |
| p-value | < 0.0001**** |  | 0.1680 |  | 0.0043** |  |

**SI Appendix, Table S2.** Statistical details for the pharmacokinetic parameters of morphine and M3G in the blood obtained from the NCA (Figure 2). Ordinary two-way ANOVA was used to assess the differences between the pharmacokinetic parameters reported for each group. Cmax, maximal concentration reached over the time course; AUC, area under the curve; AUMC, area under the first moment curve; MRT, mean residence time; Cl/F, clearance over bioavailability; V_dss_/F, volume of distribution at steady-state over bioavailability.

|  | Morphine | M3G | M3G/Morphine |
| --- | --- | --- | --- |
| **C<sub>max</sub>** |  |  |  |
| Interaction | F (1, 33) = 10.99<br>p = 0.0022** | F (1, 33) = 3.07<br>p = 0.089 | / |
| Treatment | F (1, 33) = 5.37<br>p = 0.027* | F (1, 33) = 2.33<br>p = 0.14 | / |
| Sex | F (1, 33) = 1.60<br>p = 0.21 | F (1, 33) = 34.42<br>p < 0.0001**** | / |
| **AUC** |  |  |  |
| Interaction | F (1, 32) = 8.54<br>p = 0.0063** | F (1, 32) = 2.79<br>p = 0.11 | F (1, 31) = 1.26<br>p = 0.27 |
| Treatment | F (1, 32) = 10.92<br>p = 0.0023** | F (1, 32) = 1.11<br>p = 0.30 | F (1, 31) = 7.44<br>p = 0.01* |
| Sex | F (1, 32) = 0.022<br>p = 0.88 | F (1, 32) = 18.20<br>p = 0.0002*** | F (1, 31) = 10.31<br>p = 0.0031** |
| **AUMC** |  |  |  |
| Interaction | F (1, 32) = 12.44<br>p = 0.0013** | F (1, 32) = 0.31<br>p = 0.58 | / |
| Treatment | F (1, 32) = 11.71<br>p = 0.0017** | F (1, 32) = 0.052<br>p = 0.82 | / |
| Sex | F (1, 32) = 6.59<br>p = 0.015* | F (1, 32) = 1.88<br>p = 0.18 | / |
| **MRT** |  |  |  |
| Interaction | F (1, 32) = 0.88<br>p = 0.36 | F (1, 32) = 0.45<br>p = 0.51 | / |
| Treatment | F (1, 32) = 0.0055<br>p = 0.94 | F (1, 32) = 0.49<br>p = 0.49 | / |
| Sex | F (1, 32) = 11.16<br>p = 0.0021** | F (1, 32) = 0.088<br>p = 0.77 | / |
| **T<sub>1/2</sub>** |  |  |  |
| Interaction | F (1, 32) = 0.88<br>p = 0.36 | / | / |
| Treatment | F (1, 32) = 0.0055<br>p = 0.94 | / | / |
| Sex | F (1, 32) = 11.16<br>p = 0.0021** | / | / |
| **Cl/F** |  |  |  |
| Interaction | F (1, 32) = 6.31<br>p = 0.017* | / | / |
| Treatment | F (1, 32) = 13.34<br>p = 0.0009*** | / | / |
| Sex | F (1, 32) = 0.14<br>p = 0.71 | / | / |
|  | **Vdss/F** |  |  |
| Interaction | F (1, 32) = 0.81<br>p = 0.37 | / | / |
| Treatment | F (1, 32) = 4.97<br>p = 0.033* | / | / |
| Sex | F (1, 32) = 3.11<br>p = 0.087 | / | / |

**SI Appendix, Table S3.**
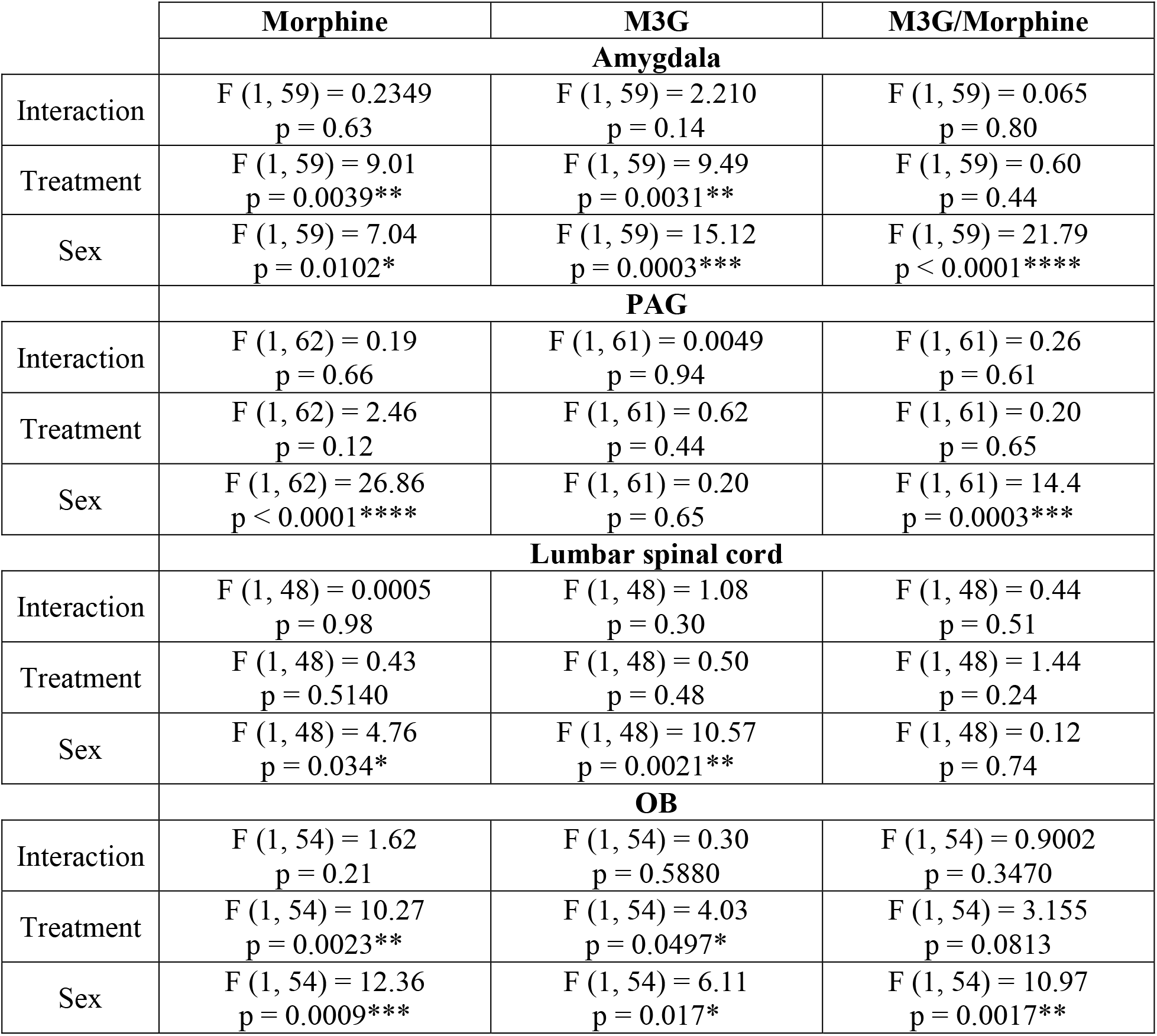
Statistical details for the quantification of morphine and M3G in the brain and lumbar spinal cord (Figure 3). Ordinary two-way ANOVA was used to assess the differences in morphine and M3G quantities between the groups.

|  | **Morphine** | **M3G** | **M3G/Morphine** |
| --- | --- | --- | --- |
|  | **Amygdala** |  |  |
| Interaction | F (1, 59) = 0.2349<br>p = 0.63 | F (1, 59) = 2.210<br>p = 0.14 | F (1, 59) = 0.065<br>p = 0.80 |
| Treatment | F (1, 59) = 9.01<br>p = 0.0039** | F (1, 59) = 9.49<br>p = 0.0031** | F (1, 59) = 0.60<br>p = 0.44 |
| Sex | F (1, 59) = 7.04<br>p = 0.0102* | F (1, 59) = 15.12<br>p = 0.0003*** | F (1, 59) = 21.79<br>p < 0.0001**** |
|  | **PAG** |  |  |
| Interaction | F (1, 62) = 0.19<br>p = 0.66 | F (1, 61) = 0.0049<br>p = 0.94 | F (1, 61) = 0.26<br>p = 0.61 |
| Treatment | F (1, 62) = 2.46<br>p = 0.12 | F (1, 61) = 0.62<br>p = 0.44 | F (1, 61) = 0.20<br>p = 0.65 |
| Sex | F (1, 62) = 26.86<br>p < 0.0001**** | F (1, 61) = 0.20<br>p = 0.65 | F (1, 61) = 14.4<br>p = 0.0003*** |
|  | **Lumbar spinal cord** |  |  |
| Interaction | F (1, 48) = 0.0005<br>p = 0.98 | F (1, 48) = 1.08<br>p = 0.30 | F (1, 48) = 0.44<br>p = 0.51 |
| Treatment | F (1, 48) = 0.43<br>p = 0.5140 | F (1, 48) = 0.50<br>p = 0.48 | F (1, 48) = 1.44<br>p = 0.24 |
| Sex | F (1, 48) = 4.76<br>p = 0.034* | F (1, 48) = 10.57<br>p = 0.0021** | F (1, 48) = 0.12<br>p = 0.74 |
|  | **OB** |  |  |
| Interaction | F (1, 54) = 1.62<br>p = 0.21 | F (1, 54) = 0.30<br>p = 0.5880 | F (1, 54) = 0.9002<br>p = 0.3470 |
| Treatment | F (1, 54) = 10.27<br>p = 0.0023** | F (1, 54) = 4.03<br>p = 0.0497* | F (1, 54) = 3.155<br>p = 0.0813 |
| Sex | F (1, 54) = 12.36<br>p = 0.0009*** | F (1, 54) = 6.11<br>p = 0.017* | F (1, 54) = 10.97<br>p = 0.0017** |

**SI Appendix, Table S4.**
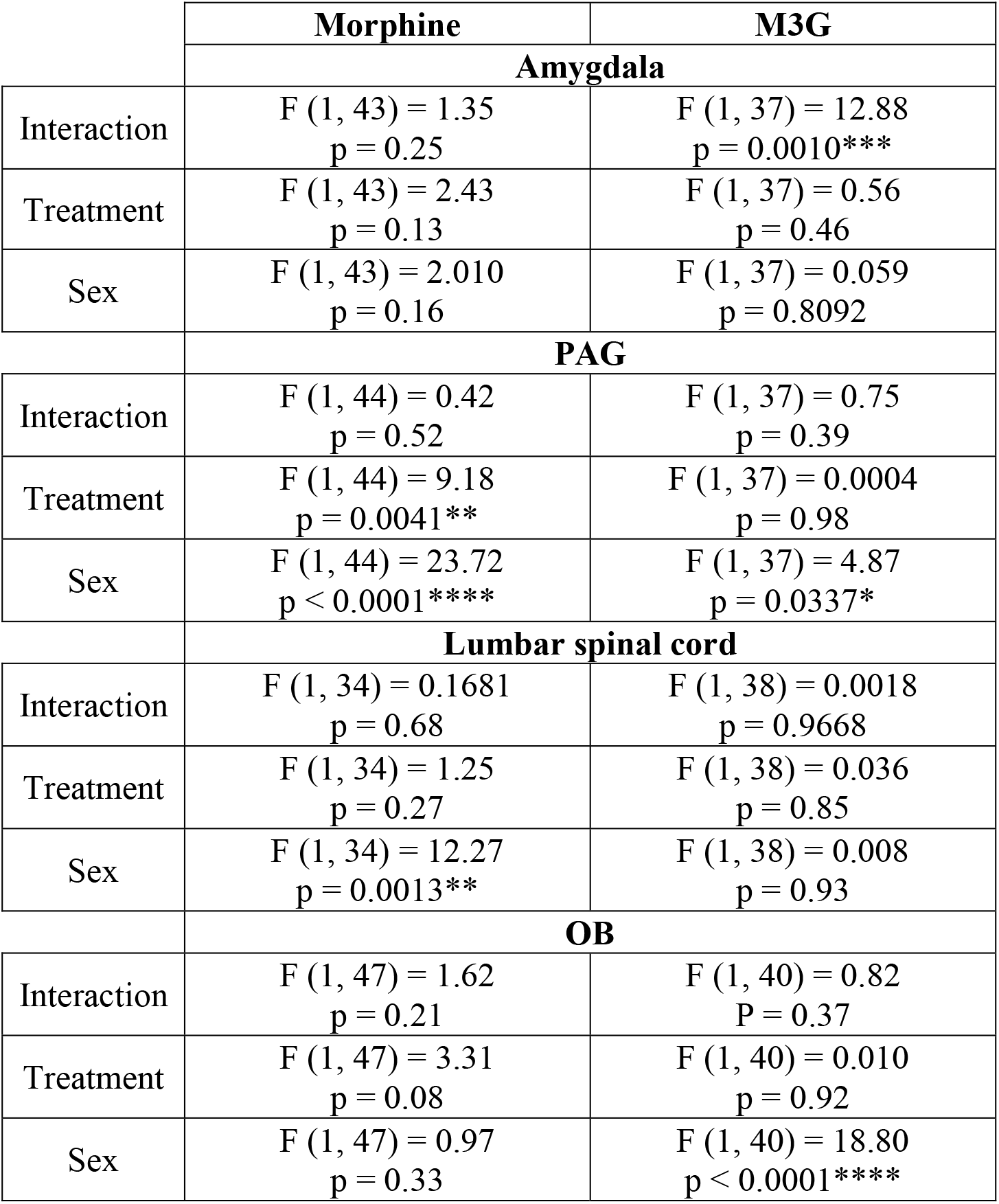
Statistical details for morphine and M3G brain/blood ratio (Figure 4). Ordinary two-way ANOVA was used to assess the differences in morphine and M3G brain/blood ratio between the groups.

**SI Appendix, Table S5.** Statistical details for M3G BBB permeability and central metabolism of morphine (Figure 4). Linear regressions were applied and analyzed through a nested-model comparison with the extra sum-of-squares F-test to compare the M3G BBB permeability between males and females, evaluate whether a significant morphine central metabolism is observed (*i.e.* comparison of the M3G brain/blood ratio models obtained after an injection of morphine or M3G) and to compare this central metabolism between males and females.

|  | BBB permeability | Morphine central metabolism :<br>Injection M3G vs Morphine | Sex differences in morphine central metabolism |
| --- | --- | --- | --- |
| **Amygdala** |  |  |  |
| Males | / | F (1, 22) = 1.61<br>p = 0.22 | / |
| Females | / | F (1, 24) = 6.74<br>p = 0.016* | / |
| M vs F | F (2, 24) = 0.046<br>p = 0.96 | / | F (1, 20) = 12.84<br>p = 0.0019** |
| **PAG** |  |  |  |
| Males | / | F (1, 25) = 11.53<br>p = 0.0023** | / |
| Females | / | F (1, 23) = 10.25<br>p = 0.0040** | / |
| M vs F | F (2, 24) = 0.41<br>p = 0.67 | / | F (1, 22) = 0.052<br>p = 0.82 |
| **ISC** |  |  |  |
| Males | / | F (1, 25) = 1.88<br>p = 0.18 | / |
| Females | / | F (1, 25) = 3.02<br>p = 0.095 | / |
| M vs F | F (2, 25) = 0.059<br>p = 0.94 | / | F (1, 23) = 0.12<br>p = 0.73 |
| **OB** |  |  |  |
| Males | / | F (2, 24) = 6.27<br>p = 0.0064** | / |
| Females | / | F (2, 23) = 2.79<br>p = 0.082 | / |
| M vs F | F (2, 25) = 0.66<br>p = 0.53 | / | F (1, 24) = 8.52<br>p = 0.0075** |

**SI Appendix, Table S6.** LC-MS/MS conditions. LC and MS/MS conditions for the purification, detection and quantification of morphine and M3G and their respective heavy-tagged counterparts. The flow rate was set at 90 μl/min on a ZORBAX SB-C18 column (150 x 1mm, 3.5μm).

|  | **Mobile phase** |  |  |
| --- | --- | --- | --- |
|  | **ACN** | **H<sub>2</sub>O** | **Formic acid** |
| **Mobile phase A** | 1% | 98.9% | 0.1% |
| **Mobile phase B** | 99.9% | 0 | 0.1% |

| **HPLC gradient** |  |  |  |  |  |  |
| --- | --- | --- | --- | --- | --- | --- |
| **Time (min)** | 0 | 2.5 | 3.5 | 4.5 | 5.0 | 8 |
| **% B mobile phase** | 0 | 0 | 98 | 98 | 0 | 0 |

| **MS parameters** |  |
| --- | --- |
| **Mode** | positive |
| **Spray voltage** | 3,500 V |
| **Nebulizer gas** | Nitrogen |
| **Desolvation (nitrogen) sheath gas** | 18 Arb |
| **Aux gas** | 7 Arb |
| **Ion transfer tube temperature** | 297°C |
| **Vaporizer temperature** | 131°C |
| **Q1 and Q3 resolutions** | 0.7 FWHM |
| **Collision gas (CID, argon) pressure** | 2 mTorr |

| **MS ionization, selection, fragmentation and identification parameters** |  |  |  |  |  |  |
| --- | --- | --- | --- | --- | --- | --- |
| **Compound** | **Polarity** | **Precursor (m/z)** | **Product (m/z)** | **Ion product type** | **Collision Energy (V)** | **RF Lens (V)** |
| **Morphine** | Positive | 285.98 | 201.11 | Quantification | 26.23 | 183 |
| **Morphine** | Positive | 285.98 | 165.36 | Qualification | 40.89 | 183 |
| **Morphine** | Positive | 285.98 | 181.06 | Qualification | 36.24 | 183 |
| **D3-morphine** | Positive | 289.1 | 201.06 | Quantification | 26.48 | 178 |
| **D3-morphine** | Positive | 289.1 | 153.13 | Qualification | 43.16 | 178 |
| **D3-morphine** | Positive | 289.1 | 165.04 | Qualification | 39.02 | 178 |
| **M3G** | Positive | 462.19 | 286.11 | Quantification | 30.02 | 276 |
| **D3-M3G** | Positive | 465.19 | 289.17 | Quantification | 29.92 | 242 |

**SI Appendix, Fig. S1.**
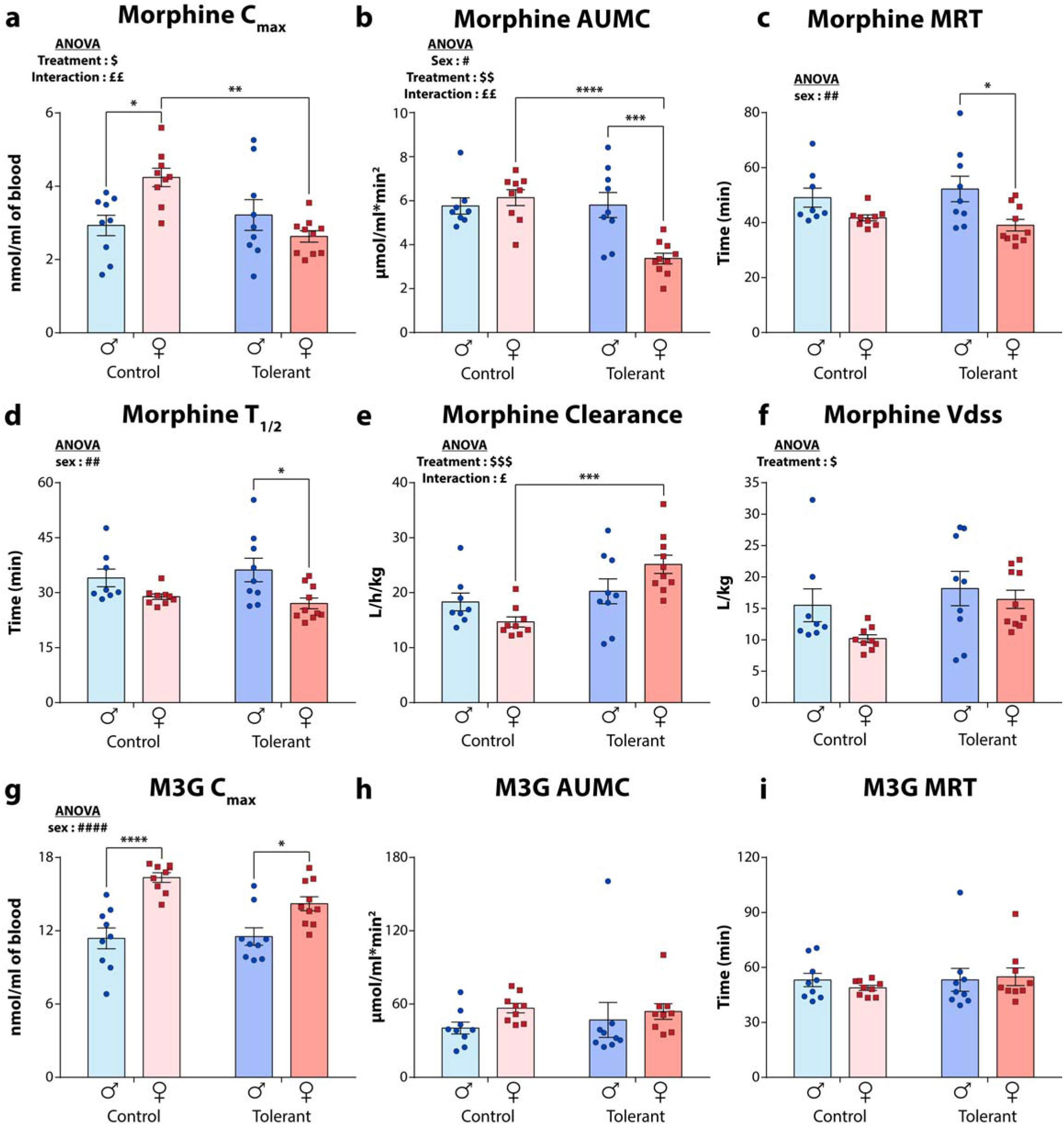
Pharmacokinetic parameters for morphine and M3G obtained from the NCA. Values of parameters obtained for (**a**) Morphine C_max_, (**b**) morphine AUMC, (**c**) morphine MRT, (**d**) morphine half-life, (**e**) morphine clearance, (**f**) morphine V_dss_, (**g**) M3G C_max_, (**h**) M3G AUMC and (**i**) M3G MRT. Values are expressed as means ± SEM, *n =* 8-10. Ordinary two-way ANOVA followed by Tukey’s multiple comparisons test was applied. Sex: #, *P*<0.05; ##, *P*<0.01; ###, *P*<0.001; ####, *P*<0.0001. Treatment: $, *P*<0.05; $$, *P*<0.01; $$$, *P*<0.001. Interaction: £, *P*<0.05; ££, *P*<0.01. *, *P*<0.05; **, *P*<0.01; ***, *P*<0.001; ****, *P*<0.0001. Males are represented as blue circle dots and females as red square dots.

**SI Appendix, Fig. S2.**
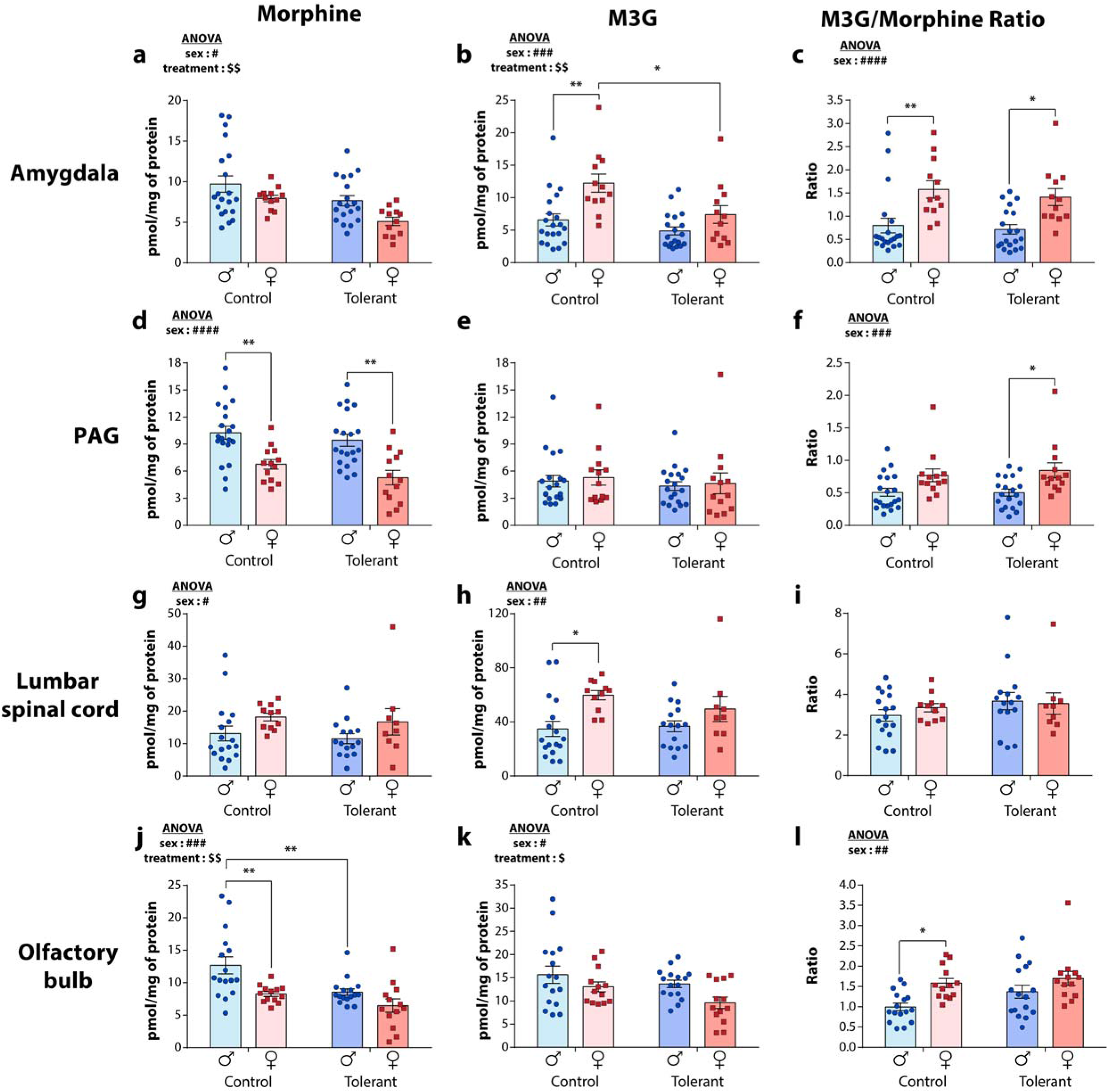
Quantities of morphine and M3G in the different brain areas and lumbar spinal cord of male and female control and tolerant mice. Levels of (**a**) morphine, (**b**) M3G and (**c**) M3G/morphine metabolic ratios present in the amygdala. Levels of (**d**) morphine, (**e**) M3G and (**f**) M3G/morphine metabolic ratios present in the PAG. Levels of (**g**) morphine, (**h**) M3G and (**i**) M3G/morphine metabolic ratios present in the lSC. Levels of (**j**) morphine, (**k**) M3G and (**l**) M3G/morphine metabolic ratios present in the OB. Values are expressed as means ± SEM, *n =* 9-20. Ordinary two-way ANOVA followed by Tukey’s multiple comparisons test was applied. Sex: #, *P*<0.05; ##, *P*<0.01; ###, *P*<0.001; ####, *P*<0.0001. Treatment: $, *P*<0.05; $$, *P*<0.01. *, *P*<0.05; **, *P*<0.01. Males are represented as blue circle dots and females as red square dots.

**SI Appendix, Fig. S3.**
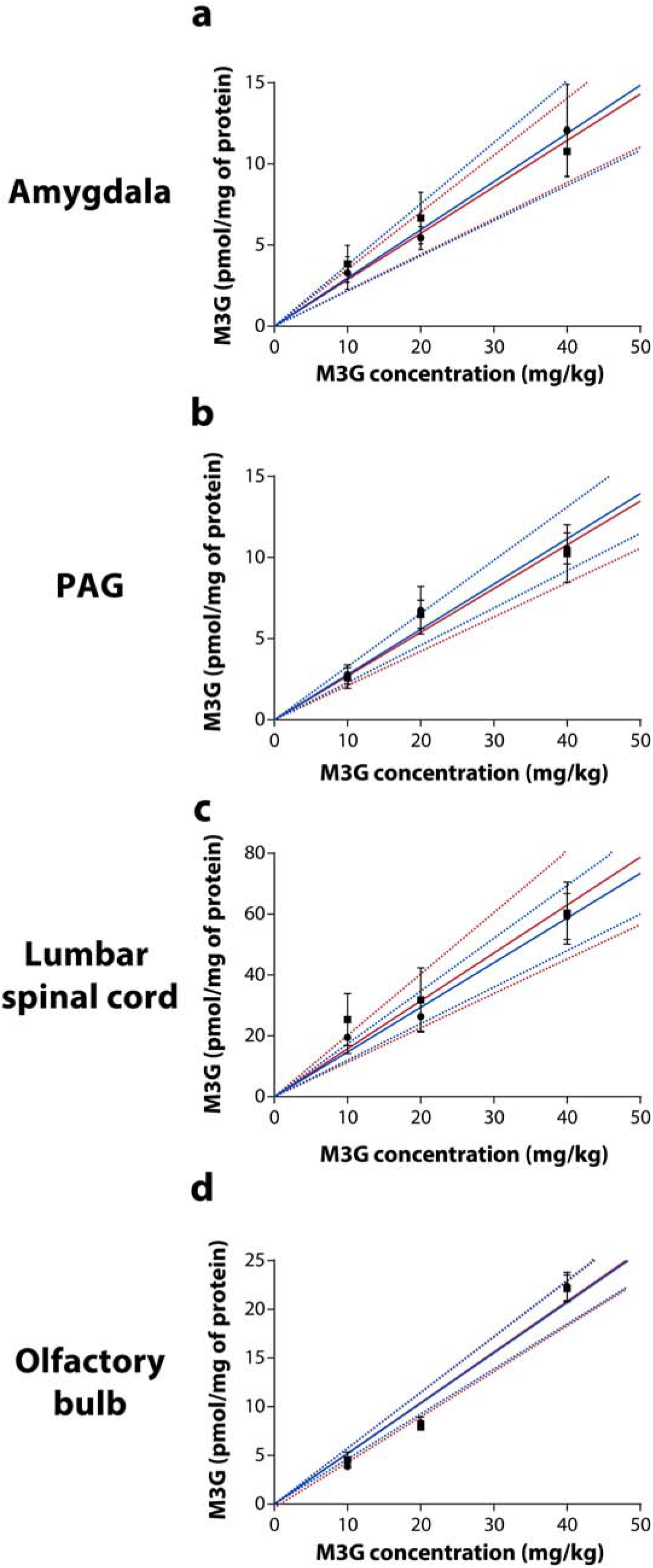
Quantities of M3G found in the different brain areas and lumbar spinal cord of male and female naïve mice following i.p. injection of increasing concentration of M3G. Levels of M3G found in (**a**) the amygdala, (**b**) the PAG, (**c**) the lSC and (**d**) the OB following i.p. injection of 10, 20 and 40 mg/kg of M3G. The blue and red lines represent linear regressions of the M3G quantities found in the brain area and lSC as a function of increasing concentration of M3G injected in males and females, respectively. Values are expressed as means ± SEM, *n =* 3-5 per sex per concentration. Males are represented as circle dots and females as square dots.

## Notes

### Competing Interest Statement

The authors have declared no competing interest.

## REFERENCES

1. Trescot AM, Datta S, Lee M, & Hansen H (2008) Opioid pharmacology. Pain Physician 11(2 Suppl):S133–153.

2. Fullerton EF, Doyle HH, & Murphy AZ (2018) Impact of sex on pain and opioid analgesia: a review. Curr Opin Behav Sci 23:183–190.

3. Cepeda MS & Carr DB (2003) Women experience more pain and require more morphine than men to achieve a similar degree of analgesia. Anesth Analg 97(5):1464–1468.

4. Comer SD, et al. (2010) Evaluation of potential sex differences in the subjective and analgesic effects of morphine in normal, healthy volunteers. Psychopharmacology (Berl) 208(1):45–55.

5. Paller CJ, Campbell CM, Edwards RR, & Dobs AS (2009) Sex-based differences in pain perception and treatment. Pain Med 10(2):289–299.

6. Eisenstein TK (2019) The Role of Opioid Receptors in Immune System Function. Front Immunol 10:2904.

7. Laux-Biehlmann A, Mouheiche J, Veriepe J, & Goumon Y (2013) Endogenous morphine and its metabolites in mammals: history, synthesis, localization and perspectives. Neuroscience 233:95–117.

8. Stone AN, Mackenzie PI, Galetin A, Houston JB, & Miners JO (2003) Isoform selectivity and kinetics of morphine 3- and 6-glucuronidation by human udp-glucuronosyltransferases: evidence for atypical glucuronidation kinetics by UGT2B7. Drug Metab Dispos 31(9):1086–1089.

9. Kurita A, et al. (2017) Comprehensive Characterization of Mouse UDP-Glucuronosyltransferase (Ugt) Belonging to the Ugt2b Subfamily: Identification of Ugt2b36 as the Predominant Isoform Involved in Morphine Glucuronidation. J Pharmacol Exp Ther 361(2):199–208.

10. Lotsch J & Geisslinger G (2001) Morphine-6-glucuronide: an analgesic of the future? Clin Pharmacokinet 40(7):485–499.

11. Lewis SS, et al. (2010) Evidence that intrathecal morphine-3-glucuronide may cause pain enhancement via toll-like receptor 4/MD-2 and interleukin-1beta. Neuroscience 165(2):569–583.

12. Smith MT, Watt JA, & Cramond T (1990) Morphine-3-glucuronide--a potent antagonist of morphine analgesia. Life Sci 47(6):579–585.

13. Weinsanto I, et al. (2018) Stable isotope-labelled morphine to study in vivo central and peripheral morphine glucuronidation and brain transport in tolerant mice. Br J Pharmacol 175(19):3844–3856.

14. Williams JT, et al. (2013) Regulation of mu-opioid receptors: desensitization, phosphorylation, internalization, and tolerance. Pharmacol Rev 65(1):223–254.

15. Eidson LN, Inoue K, Young LJ, Tansey MG, & Murphy AZ (2017) Toll-like Receptor 4 Mediates Morphine-Induced Neuroinflammation and Tolerance via Soluble Tumor Necrosis Factor Signaling. Neuropsychopharmacology 42(3):661–670.

16. Due MR, et al. (2012) Neuroexcitatory effects of morphine-3-glucuronide are dependent on Toll-like receptor 4 signaling. J Neuroinflammation 9(1):200.

17. Bai L, et al. (2014) Toll-like receptor 4-mediated nuclear factor-kappaB activation in spinal cord contributes to chronic morphine-induced analgesic tolerance and hyperalgesia in rats. Neurosci Bull 30(6):936–948.

18. Mattioli TA, et al. (2014) Toll-like receptor 4 mutant and null mice retain morphine-induced tolerance, hyperalgesia, and physical dependence. PLoS One 9(5):e97361.

19. Roeckel LA, et al. (2017) Morphine-induced hyperalgesia involves mu opioid receptors and the metabolite morphine-3-glucuronide. Sci Rep 7(1):10406.

20. Kest B, Wilson SG, & Mogil JS (1999) Sex differences in supraspinal morphine analgesia are dependent on genotype. J Pharmacol Exp Ther 289(3):1370–1375.

21. Kest B, Palmese C, & Hopkins E (2000) A comparison of morphine analgesic tolerance in male and female mice. Brain Res 879(1-2):17–22.

22. Mogil JS (2012) Sex differences in pain and pain inhibition: multiple explanations of a controversial phenomenon. Nat Rev Neurosci 13(12):859–866.

23. Sarton E, et al. (2000) Sex differences in morphine analgesia: an experimental study in healthy volunteers. Anesthesiology 93(5):1245–1254; discussion 1246A.

24. Craft RM (2003) Sex differences in opioid analgesia: "from mouse to man". Clin J Pain 19(3):175–186.

25. Kest B, Sarton E, & Dahan A (2000) Gender differences in opioid-mediated analgesia: animal and human studies. Anesthesiology 93(2):539–547.

26. Cicero TJ, Nock B, O'Connor L, & Meyer ER (2002) Role of steroids in sex differences in morphine-induced analgesia: activational and organizational effects. J Pharmacol Exp Ther 300(2):695–701.

27. Loyd DR, Wang X, & Murphy AZ (2008) Sex differences in micro-opioid receptor expression in the rat midbrain periaqueductal gray are essential for eliciting sex differences in morphine analgesia. J Neurosci 28(52):14007–14017.

28. Loyd DR & Murphy AZ (2014) The neuroanatomy of sexual dimorphism in opioid analgesia. Exp Neurol 259:57–63.

29. Doyle HH, Eidson LN, Sinkiewicz DM, & Murphy AZ (2017) Sex Differences in Microglia Activity within the Periaqueductal Gray of the Rat: A Potential Mechanism Driving the Dimorphic Effects of Morphine. J Neurosci 37(12):3202–3214.

30. Soldin OP & Mattison DR (2009) Sex differences in pharmacokinetics and pharmacodynamics. Clin Pharmacokinet 48(3):143–157.

31. South SM, Edwards SR, & Smith MT (2009) Antinociception versus serum concentration relationships following acute administration of intravenous morphine in male and female Sprague-Dawley rats: differences between the tail flick and hot plate nociceptive tests. Clin Exp Pharmacol Physiol 36(1):20–28.

32. Chen J, et al. (2017) Profiles and Gender-Specifics of UDP-Glucuronosyltransferases and Sulfotransferases Expressions in the Major Metabolic Organs of Wild-Type and Efflux Transporter Knockout FVB Mice. Mol Pharm 14(9):2967–2976.

33. Rush GF, Newton JF, & Hook JB (1983) Sex differences in the excretion of glucuronide conjugates: the role of intrarenal glucuronidation. J Pharmacol Exp Ther 227(3):658–662.

34. Bond JA, Medinsky MA, Dent JG, & Rickert DE (1981) Sex-dependent metabolism and biliary excretion of [2,4-14C] dinitrotoluene in isolated perfused rat livers. J Pharmacol Exp Ther 219(3):598–603.

35. Baker L & Ratka A (2002) Sex-specific differences in levels of morphine, morphine-3-glucuronide, and morphine antinociception in rats. Pain 95(1-2):65–74.

36. Mackenzie PI, et al. (2003) Regulation of UDP glucuronosyltransferase genes. Curr Drug Metab 4(3):249–257.

37. Yamada H, et al. (2003) Formation of highly analgesic morphine-6-glucuronide following physiologic concentration of morphine in human brain. J Toxicol Sci 28(5):395–401.

38. Bickel U, Schumacher OP, Kang YS, & Voigt K (1996) Poor permeability of morphine 3-glucuronide and morphine 6-glucuronide through the blood-brain barrier in the rat. J Pharmacol Exp Ther 278(1):107–113.

39. Murphey LJ & Olsen GD (1994) Diffusion of morphine-6-beta-D-glucuronide into the neonatal guinea pig brain during drug-induced respiratory depression. J Pharmacol Exp Ther 271(1):118–124.

40. Handal M, Grung M, Skurtveit S, Ripel A, & Morland J (2002) Pharmacokinetic differences of morphine and morphine-glucuronides are reflected in locomotor activity. Pharmacol Biochem Behav 73(4):883–892.

41. Bartlett SE, Cramond T, & Smith MT (1994) The excitatory effects of morphine-3-glucuronide are attenuated by LY274614, a competitive NMDA receptor antagonist, and by midazolam, an agonist at the benzodiazepine site on the GABAA receptor complex. Life Sci 54(10):687–694.

42. Peckham EM & Traynor JR (2006) Comparison of the antinociceptive response to morphine and morphine-like compounds in male and female Sprague-Dawley rats. J Pharmacol Exp Ther 316(3):1195–1201.

43. Swartjes M, et al. (2012) Morphine Induces Hyperalgesia without Involvement of mu-Opioid Receptor or Morphine-3-glucuronide. Molecular Medicine 18(9):1320–1326.

44. Penson RT, et al. (2000) Randomized placebo-controlled trial of the activity of the morphine glucuronides. Clin Pharmacol Ther 68(6):667–676.

45. Barjavel MJ, Scherrmann JM, & Bhargava HN (1995) Relationship between morphine analgesia and cortical extracellular fluid levels of morphine and its metabolites in the rat: a microdialysis study. Br J Pharmacol 116(8):3205–3210.

46. Craft RM, Stratmann JA, Bartok RE, Walpole TI, & King SJ (1999) Sex differences in development of morphine tolerance and dependence in the rat. Psychopharmacology (Berl) 143(1):1–7.

47. Aloisi AM, et al. (2010) Aromatase and 5-alpha reductase gene expression: modulation by pain and morphine treatment in male rats. Mol Pain 6:69.

48. Richner M, Jager SB, Siupka P, & Vaegter CB (2017) Hydraulic Extrusion of the Spinal Cord and Isolation of Dorsal Root Ganglia in Rodents. J Vis Exp (119).

49. Ho YS, Liu RH, Nichols AW, & Kumar SD (1990) Isotopic Analog as the Internal Standard for Quantitative-Determination - Evaluation of Mass-Spectra of Commonly Abused Drugs and Their Deuterated Analogs. Journal of Forensic Sciences 35(1):123–132.

50. Zhang Y, Huo M, Zhou J, & Xie S (2010) PKSolver: An add-in program for pharmacokinetic and pharmacodynamic data analysis in Microsoft Excel. Comput Methods Programs Biomed 99(3):306–314.

